# Graded BMP signals modulate yellow and red color in fishes impacting adult pigment pattern and behavior

**DOI:** 10.64898/2025.12.04.692427

**Authors:** Delai Huang, Pietro L. H. de Mello, Tiffany Liu, Yu Liu, Emaan H. Kapadia, Yipeng Liang, Jianguo Lu, Joseph C Corbo, David M Parichy

## Abstract

Among the most interesting adult traits are those with roles in animal communication. Yet developmental mechanisms by which genes drive cell behaviors in building the final forms of such traits are rarely known. In this context, pigmentation is useful because colors and patterns often provide signals in mate choice, predation avoidance and other behaviors and pigmentation is unusually accessible to observation and manipulation. Here we focus on some of the most prominent signaling colors—red, orange and yellow—and show how BMP signaling at the cellular level allows for a very different kind of signal at the organismal level. Using pearl danio, *Danio albolineatus*, we find that spatially and temporally graded BMP signals promote development of yellow/orange xanthophores over red erythrophores in the fin of this species and a distantly related minnow, *Tanichthys albonubes*, and that conserved mechanisms, involving BMP co-receptor Rgmb, regulate differentiation of other pigment cell types in corresponding locations of zebrafish, *D. rerio*. We further use mutants of *D. albolineatus* with more red or more yellow cells than wild-type to demonstrate female responsiveness to carotenoid-based color differences between males in shoaling preference assays, and we show the existence of polygenic standing variation for this pigmentary trait. Our findings illustrate a chain of function spanning hierarchical levels and provide a deeper understanding of pigmentary form and function and its evolution.

**HIGHLIGHTS:**

- Fate specification of alternative red or yellow pigment cell types in the fin of a minnow, pearl danio, depends on thresholds and gradients in BMP signaling during fin outgrowth.
- Genetic loss of BMP signaling leads to production of red over yellow carotenoids with resulting “super-red” fish preferred by females in shoaling assays.
- BMP-dependence of pigmentary traits is conserved across species with standing, polygenic variation for fin pattern and color in pearl danio.

## RESULTS AND DISCUSSION

A wide variety of morphological features are adapted for signaling between individuals: the peacock’s ornate plumage that handicaps survival and so tells of underlying genetic quality, the enlarged claws of fiddler crabs that attract females and threaten rivals, and the antlers of male deer that indicate dominance and can be used to demonstrate social status when needed. The ethological context of such signals are often understood^1,2^. Occasionally, segregating alleles that modulate signal production are known. But how gene activities—from fixed alleles and segregating variants—are translated through morphogenesis and differentiation of cells into particular morphological outcomes remains largely mysterious, despite the importance of such information for understanding diversity of form and function in the natural world.

Among the many roles for integumentary pigmentation, signaling is especially common and important^3–5^. Yellow, orange and red colors often function in mate choice and warning coloration and frequently depend on carotenoids that are dietarily acquired but endogenously processed and localized^6–8^. In ectothermic vertebrates specialized chromatophores of the neural crest lineage concentrate and display these pigments: red erythrophores and yellow to orange xanthophores^9^. To uncover molecular and cellular bases for the development and arrangements of these cells we focused on minnows of the family Cyprinidae, which have diverse colors and color patterns, and, in the genus *Danio*, a phylogenetic proximity to zebrafish that makes them convenient for mechanistic and behavioral studies^10–12^.

To identify factors responsible for color and pattern we reasoned that genetic pathways required for differentiating erythrophores and xanthophores from a common progenitor^13^ could still be evident in the transcriptomes of fully pigmented cells. In pearl danio, *D. albolineatus*, xanthophores and erythrophores are intermingled on the body yet segregated to discrete regions in the fin, with erythrophores proximally, xanthophores further distally, and a region devoid of either cell, yet containing black melanophores, at the distal tip^14^ (**Figure 1A,B**). We isolated erythrophores and xanthophores by dissecting corresponding regions of fins and then sorting for fluorescence of transgenic reporter *aox5:nucEos*+, expressed by both types. Transcriptomic comparison of cells by bulk RNA-sequencing revealed several genes associated with BMP signaling expressed at markedly higher levels in xanthophores than erythrophores (**Figure 1C**; Table S1).

**Figure 1.**
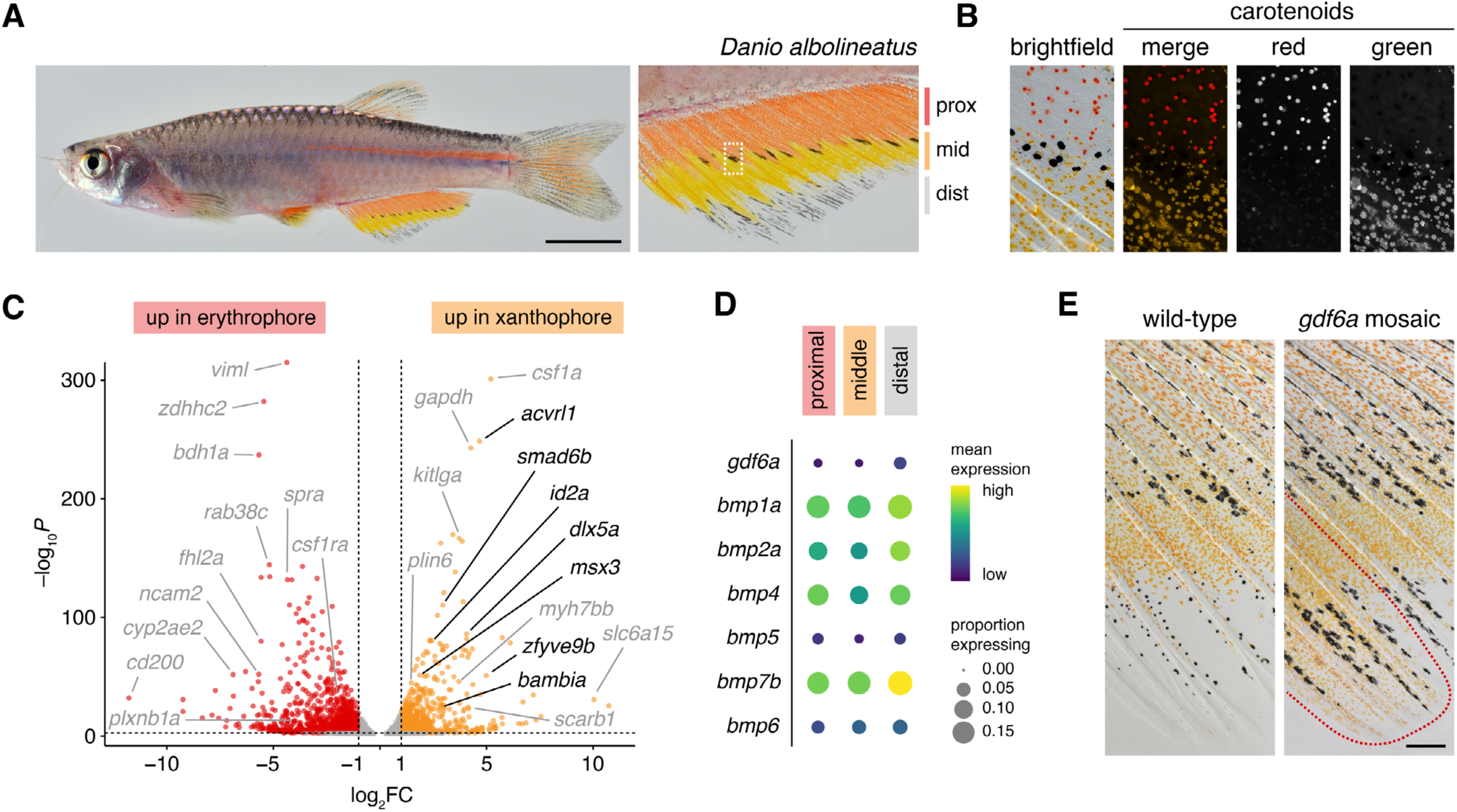
Pattern domains of erythrophores and xanthophores depend on BMP signals. (A) Wild-type male *D. albolineatus* with red erythrophores and yellow/orange xanthophores, intermingled on the body but spatially separated into proximal and distal regions in the fin; a few melanophores occur in the middle where erythrophores and xanthophores meet. (B) Boxed region in A after contracting carotenoid vesicles towards cell centers with epinephrine. Red carotenoids of erythrophores autofluorescence in the red channel whereas yellow/orange carotenoids of xanthophores autofluorescence in the green channel, here pseudocolored orange. (C) Several BMP signaling or response genes were expressed more highly in xanthophores than erythrophores (shown in black). Additional genes with documented pigmentary functions, potential impacts on morphogenesis, or possible utility as markers due to expression differences are in grey. (D) Transcripts of several BMP ligand genes were biased distally in their abundance (distal vs. proximal and middle; all *q*<0.05). (E) Mosaic loss-of-function for *gdf6a* expanded territories occupied by xanthophores towards the distal fin margin. Clones of cells extend proximodistally in fins^13,51^; dotted line here indicates location of *gdfd6a-/-* cells. Scale bars, 5 mm (A), 250 *µ*m (E).

BMPs are well-known morphogens^15–17^, driving different cellular responses at different levels of signal strength and duration while also exhibiting their own context-specific activities^18–21^. BMPs also function in outgrowth and regeneration of fish fins^22,23^. We therefore hypothesized that spatially or temporally graded BMP signals specify xanthophore vs. erythrophore fates. If so, genes encoding BMP or related ligands might be expressed differentially along the fin proximodistal axis. This prediction was borne-out by transcriptomic comparisons of single cells in the tissue environment of developing pigment cells (**Figure 1D**; **Figure S1A–C**; Tables S02, S03), with particularly notable expression differences for *gdf6a* and *bmp7b* within a population of presumptive osteoblast progenitor cells marked by high levels of expression for genes encoding collagens (types I, V, XII), Periostin, Parathyroid hormone 1 receptor and Fgf receptor (**Figure S1D**).

To determine if xanthophores, erythrophores or their progenitors are competent to receive BMP signals we separately assayed expression of BMP receptors by single cell RNA-sequencing of *aox5*:nEos+ cells at an earlier stage, as their type-specific pigmentary phenotypes were being acquired. These analyses revealed several BMP receptor genes—some expressed differentially between cell types—as well as modules of gene expression associated with xanthophore and erythrophore differentiation more generally (**Figure S1E–H;** Tables S04, S05). As compared to xanthophores, early erythrophores also expressed higher levels of BMP signaling inhibitor gene *fstl1b* (**Figure S1E**) whereas later erythrophores expressed higher levels of *fstl1b* and *fstb* (log_2_FC=1.2,1.5, *q*≤1.7E-6; Table S1) suggesting diminished BMP signal reception may accompany differentiation.

Transcriptomic analyses thus pointed to differential BMP gene expression in the tissue environment, competency of xanthophores and progenitors to receive such signals, and inhibitory factors potentially attenuating signal transmission to erythrophores. As a first step in testing roles for BMP signaling in the erythrophore-xanthophore lineage we targeted the ligand gene *gdf6a*, which functions in other pigmentary contexts and allows for mutant phenotypes that are semi-viable into adult stages^24,25^. Fish genetically mosaic for CRISPR/Cas9 induced mutations survived to post-embryonic stages and developed xanthophores along the distal fin margin where these cells are normally absent (**Figure 1E**).

Changes in xanthophore distributions in *gdf6a* mosaics along with anatomical and temporal changes in BMP pathway gene expression suggested that different levels of BMP signaling might be experienced by erythrophores and xanthophores. Indeed, erythrophores exhibited little to no detectable pathway activity as inferred from immunohistochemical staining for pSmad whereas xanthophores exhibited high levels of activity (**Figure 2A**). Quantitative assessment confirmed a distal bias of anti-pSmad immunoreactivity at stages after cell-type specific colors were evident, yet proximodistal variation in staining intensity was not evident among earlier stage progenitors of nearly uniform color (**Figure S2**).

**Figure 2.**
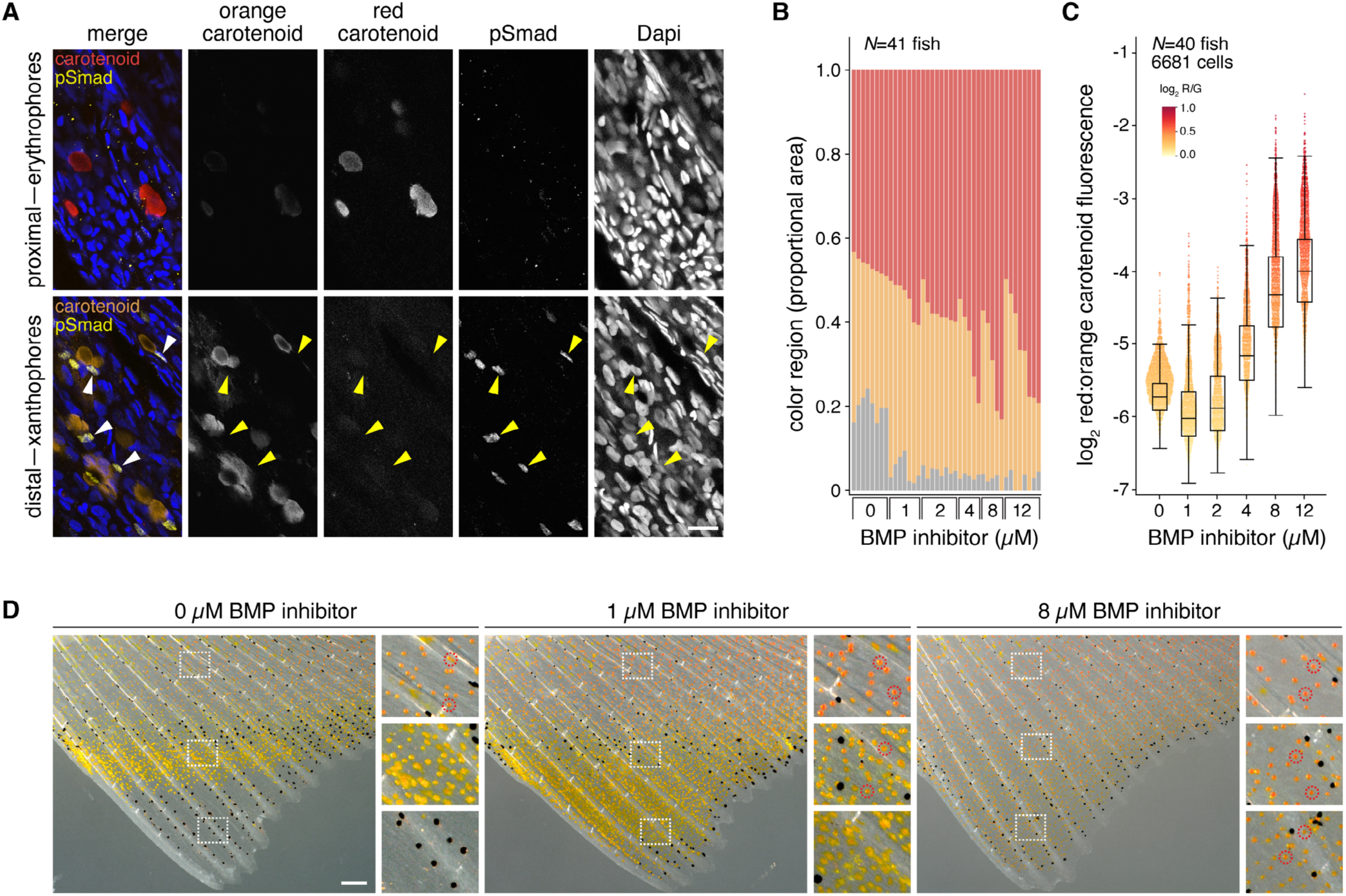
BMP signals specify xanthophores over erythrophores. (A) pSmad immunoreactivity was not evident proximally where erythrophores are located but was detected distally in nuclei of xanthophores. Different channels for detection of cell-type specific pigments are pseudocolored to match cell appearance in brightfield (red ketocarotenoids of erythrophores detected in red and shown in red; yellow carotenoids of xanthophores detected in green and shown in orange). Staining for pSmad (yellow in merge) coincided with locations of Dapi-stained nuclei and was adjacent to accumulations of pigment vesicles mobilized towards cell centers by epinephrine treatment^13^. (B) Higher proportions of fins were covered by erythrophores than xanthophores upon treatment with BMP inhibitor. Each bar represents a single individual, ordered within treatments by extent of red coverage and showing distal regions without either cell type in grey. (C) BMP inhibition also shifted ratios of red to yellow/orange carotenoids across cells measured individually for autofluorescence signatures (as in A). (D) Representative fins showing increasingly distal positioning of erythrophores and xanthophores with BMP inhibition. A few of many example erythrophores, with red pigments, are circled in insets. Scale bars, 20 *µ*m (A), 200 *µ*m (D).

These observations suggested a model for chromatophore patterning during fin outgrowth^13^ in which low BMP signals permit erythrophore differentiation proximally, higher levels induce xanthophore differentiation in middle-to-distal regions, and highest levels prevent xanthophores from differentiating at the distal tip. To test this idea, we inhibited BMP signaling during fin outgrowth beginning just prior to erythrophore and xanthophore differentiation using LDN-193189 ^(26,27)^, which binds to BMP type I receptors and reduced pSmad immunostaining as expected (**Figure S3A**). Increasing doses of BMP inhibitor led to increased areas covered by erythrophores, which extended further towards the fin tip, as well as increased redness within individual cells as assayed by relative fluorescence of red and yellow/orange carotenoids^13^ (**Figure 2B–D**). Distal limits of xanthophore regions shifted markedly towards fin tips. At low doses, total areas covered by xanthophores tended to be greater than in controls and at high doses these areas were far more variable among individuals than observed for controls (**Figure S3B**). These outcomes are consistent with flattening a signaling gradient: regions permissive for xanthophore development could be smaller or larger than controls, depending on efficacy of inhibition across individuals and whether levels of signal fell within thresholds for xanthophore differentiation. Changes in pattern reflected alterations in locations and fates of cells differentiating from progenitors, rather than death, migration or transdifferentiation of cells present at the start of treatments (**Figure S3C,D**). Although fin melanophores normally positioned between erythrophores and xanthophores were missing in BMP-inhibitor treated fish, loss of these cells probably did not otherwise affect patterning as a *D. albolineatus* mutant^28^ that lacks fin melanophores developed erythrophore and xanthophore boundaries at their normal locations (**Figure S4A**).

Requirements for BMP signaling in patterning adult pigment cells were intriguing so we asked whether they might be generalizable to other species, cell types, and anatomical locations. To determine if BMP requirements for specifying xanthophores over erythrophores are conserved evolutionarily we examined white cloud minnow, *Tanichthys albonubes*, a distant relative of *Danio* within Cyprinidae^29^ with both types of cells in its fins. Xanthophores but not erythrophores exhibited strong pSmad immunoreactivity, as in *D. albolineatus* (**Figure 3A**). BMP inhibition likewise caused normally proximal erythrophores to differentiate widely across the fin, with xanthophores confined to distalmost regions (**Figure 3B**). A reciprocal shift was evident on the body, manifested in a dorsal shift in the positions of a light “interstripe” of yellowish iridophores^30^ and erythrophores, along with the upper boundary to a dark stripe of melanophores and blue iridophores (**Figure S4B**).

**Figure 3.**
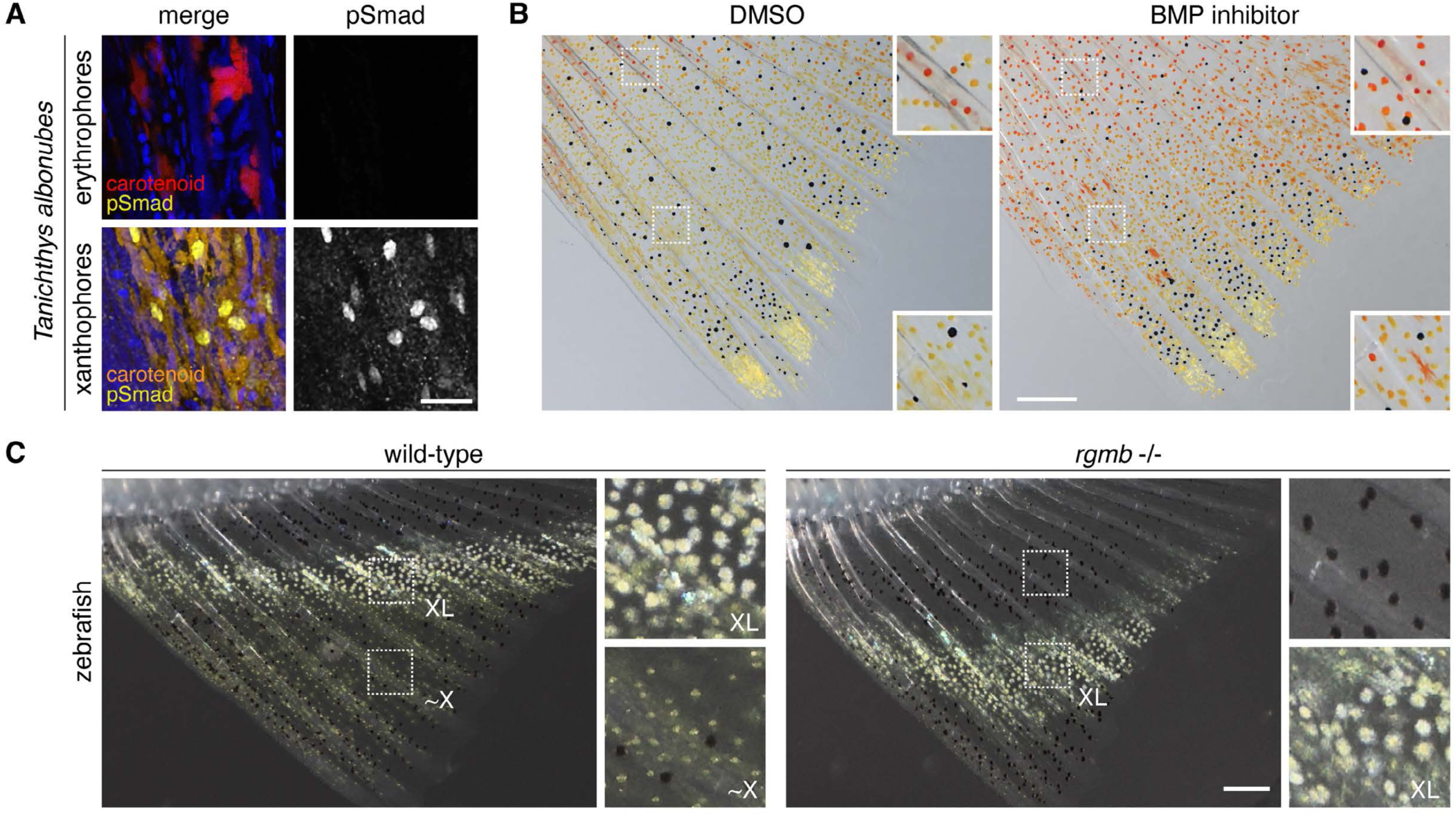
Conserved BMP signaling in pigment cell specification and pattern formation. (A) *Tanichthys albonubes* erythrophores were pSmad– whereas xanthophores were strongly pSmad+. Cytoplasmic signals in xanthophores were from pigment autofluorescence. (B) BMP inhibition led to an expanded domain of erythrophores and a distal shift of xanthophores. (C) Zebrafish early adult fin pattern with a light stripe of xantholeucophores (XL) and more distal xanthophore-like (∼X) cells thought to be progenitors of XL^31^. In the *rgmb* mutant, defective for BMP signaling, XL were displaced distally with melanophores occupying their normal location. Scale bars, 20 *µ*m (A), 200 *µ*m (B,C).

To test dependencies of still other pigment cell types and patterns on BMP signaling, we turned to zebrafish. Unlike many danios, zebrafish lack erythrophores^13^. The anal fin pattern consists of dark stripes of melanophores that alternate with light interstripes of whitish-yellow xantholeucophores that develop through a xanthophore-like state^31^ (**Figure 3C**). During pattern formation, relatively mature proximal xantholeucophores had low levels of pSmad immunostaining whereas more distal xanthophore-like cells had strong pSmad staining (**Figure S4C**). By comparison to wild-type, mutants for the non-canonical BMP coreceptor Rgmb^27,32^ had reduced pSmad signaling overall and a distal shift of mature xantholeucophores as the pattern was forming (**Figure 3C**). At later stages, proximal fin melanophores were positioned more distally, xanthophores were deficient for yellow coloration, and defects in pattern orientation were evident; alterations in body and dorsal fin pattern and color were apparent as well (**Figure S4D**). These findings point to conserved roles for BMP signals in specifying pattern across taxa and chromatophore classes.

Phenotypes of zebrafish *rgmb* mutants, and the presence of *rgmb* transcript in erythrophores and xanthophores (**Figure S1E**), led us to ask if the relative abundances of these cells might depend on Rgmb activity. Upon generating homozygous mutants for *rgmb* of *D. albolineatus* (**Figure S5A**) we found a marked increase in erythrophores, which extended to the distal tip of the fin, and a corresponding reduction in xanthophores (**Figure 4A**, left panels). Erythrophores of the mutant were more dispersed than xanthophores, as in the wild-type (**Figure 4A**, middle panels), and had large carotenoid vesicles, indistinguishable from erythrophores of wild-type fish, and larger than those of xanthophores in the wild type (**Figure S5B**). Though BMP-inhibitor treated *D. albolineatus* lacked clear defects in body pattern, a residual pattern of melanophore stripes, dorsal and ventral to an evolutionarily reduced interstripe^33^, was less organized and ultimately more diffuse in *rgmb-/-* than in the wild type (**Figure S5C** and see below).

**Figure 4.**
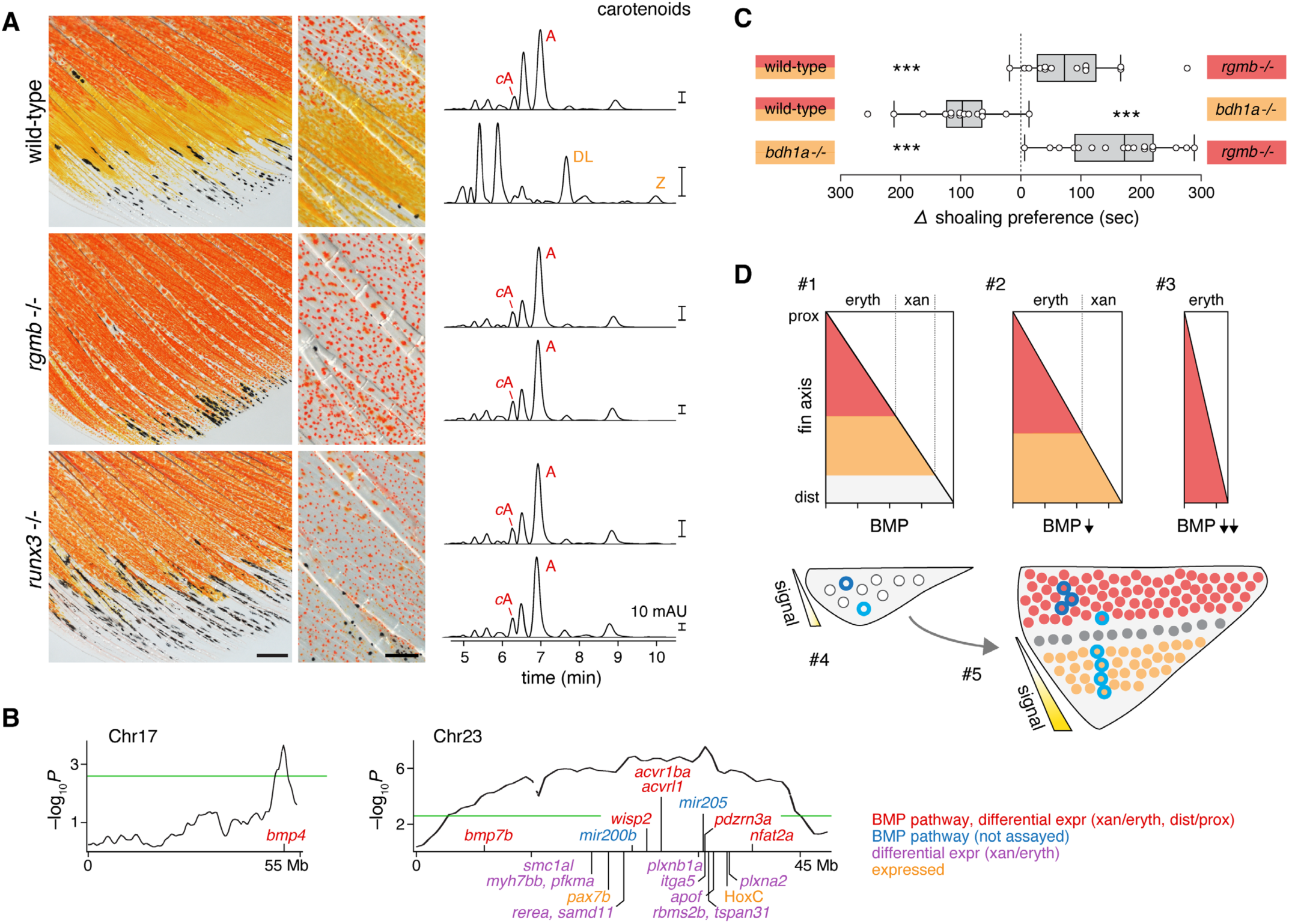
Induced and standing genetic variation for phenotype and impacts on conspecific signal response. (A) *rgmb* and *runx3* mutants were deficient for yellow/orange xanthophores. Low and high magnification images taken before and after epinephrine treatment, respectively. HPLC chromatographs (450 nm) are shown for proximal (upper) and distal (lower) fin regions from pooled fish in representative samples of each genotype. *c*A, *cis*-Astanxanthin; A, Astanxanthin; DL, Dehyro-lutein; Z, Zeaxanthin. (B) Genetic mapping for fin variation in fin pattern and color revealed multiple regions of association across the genome, including one end of Chr17 and a strong nearly chromosome-wide association with Chr23. Plots show G’ statistics with 1 Mb window (green threshold, *q*=0.05). Critical intervals included loci with alternative alleles fixed in parentals that were also differentially expressed across cell types or fin regions (red) of *D. albolineatus* (Figure 1C,D, Figure S1D,E) or were expressed in the xanthophores and erythrophores and known to function in xanthophore fate in zebrafish (*pax7b*)^52^ (C) In assays of female shoaling, *rgmb-/-* males were prefered over wild-type and both wild-type and *rgmb-/-* males were preferred over red-deficient *bdh1a-/-* males. Points show differences in total times spent by individual females in proximity to each stimulus shoal (hypothesized mean difference = 0, Wilcoxon, |*S*|≥49.5, *P*<0.001; *N*=14–17 fish tested). (D) Model for pigmentary phenotype dependence on BMP signaling and fin outgrowth. Alternative erythrophore and xanthophore fates from a common progenitor^13^ depend on different upper and lower thresholds of BMP activity (#1). Genetic mosaic or pharmacological inhibitions of signaling (*gdf6a*, LDN-193189; #2) or more severe reductions due to biallelic mutations (*rgmb-/-*, *runx3-/-*; #3) alter fate specification and proximal–distal pattern boundaries. These effects occur in the context of extensive proximodistal outgrowth from ∼0.5 mm base-to-tip when unspecified progenitors are first evident (#4) to ∼2.2 mm when earliest erythrophores and xanthophores are recognizable (#5). Prior fate mapping^13^ showed that proximal progenitors (deep blue outline) produce erythrophores whereas middle or distal progenitors (light blue outline) produce erythrophores and xanthophores or just xanthophores across the fin with some progeny being displaced distally as the fin grows. Scale bars, 500 *µ*m (left), 200 *µ*m (right).

In other contexts, transcription factor Runx3^34^ functions in concert with BMP signaling and in zebrafish it acts downstream of Rgmb in pigment cells^27^; here we found that *runx3* was expressed similarly to *rgmb* in erythrophores, xanthophores and their progenitors (**Figure S1H**). To test for a conserved role of Runx3 in fate specification downstream of Rgmb in *D. albolineatus*, we generated fish homozygous for premature termination alleles and found them to have phenotypes similar to those of *rgmb-/-*, albeit somewhat less severe (**Figure 4A**). Taken together, these findings—with those of pharmacological BMP-inhibition and *gdf6a* mosaics—suggest roles for both canonical and non-canonical BMP receptors, as well as conservation in *D. albolineatus* of a BMP/Runx3 module that affects differentiation of pigment cells.

Erythrophores and xanthophores contain different complements of carotenoids, with red colors attributable to the presence of ketocarotenoids that are produced via enzymatic addition of ketone groups to yellow carotenoids^7,13^. The overtly red appearances of *rgmb* and *runx3* mutant *D. albolineatus* implied changes in carotenoid content, which we confirmed by high performance liquid chromatography (**Figure 4A**, right panels). Examination of wild-type fish revealed the presence of multiple carotenoids in proximal fin regions, with a major peak corresponding to the ketocarotenoid astaxanthin and distinct profiles of carotenoids in distal regions, including peaks corresponding to dehydro-lutein and zeaxanthin, which confer a yellow color (**Figure S6A–C**). *rgmb* and *runx3* mutants had significantly more astaxanthan than wild-type and neither mutant had detectable levels of dehydro-lutein or zeaxanthin. Conversely, fish with mutations in *bdh1a*, which encodes an enzyme required for the formation of ketocarotenoids, lacked astaxanthin and had excess dehydro-lutein and zeaxanthin (**Figure S6C,D**; Table S6).

Having identified a pathway and specific genes in this pathway required for fin pigmentary phenotype we were curious about the broader genetic architecture of this trait and whether there might be allelic variation available for selection. In some prior studies we used a variant of pearl danio, *D.* aff. *albolineatus*^11,35^, in which individuals often have erythrophores that extend further distally and a redder appearance overall. To interrogate the genetic basis of this variation we crossed *D.* aff. *albolineatus* and *D. albolineatus*, and then crossed two F1s to generate a family of ∼300 fish that segregated variation in phenotype, from which we selected pools of extreme yellow and extreme orange-red males. Whole-genome sequencing and mapping revealed a highly significant association with phenotype across Chr23, along with weaker associations across several other chromosomes, raising the possibility of multiple contributing loci (**Figure S7**). Variant effect prediction did not point to mutations of clear functional significance yet the broad critical region on Chr23 and an additional region on Chr17 included genes expressed differentially between erythrophores and xanthophores that function in BMP/TGFβ signaling (e.g., *acvrl1*, *bmp4*, *bmp7b*) and other genes expressed by these cells with plausible relevance to differentiation or morphogenesis (**Figure 4B)**.

Finally, with a deeper understanding of developmental mechanisms and standing variation we asked if these phenotypes impact behavior, as might be anticipated given roles for red, orange or yellow colors as signals in mate choice, social aggregation, and dominance displays^36–40^. We focused on a social aggregative behavior, shoaling, important for predation avoidance, foraging efficiency and access to mates^41^. Zebrafish associate with one another based on pigmentary and other factors^42–44^ but cues used by *D. albolineatus* in shoaling with conspecifics are not known. We reasoned that isolation of extreme phenotypes beyond those of wild-type (“super-red” *rgmb* and red-deficient *bdh1a*; **Figure S8A,B**) provided an opportunity to test if these colors provide signals that impact shoaling.

Proximity to conspecifics is a necessary prelude to mate choice and *D. albolineatus* are sexually dimorphic, with males redder than females^13^, so we tested shoaling of females in binary preference assays using alternative male phenotypes. When presented with wild-type and *rgmb* mutants, females preferred to shoal with *rgmb-/-* (**Figure 4C**, **Figure S8C**). This preference could indicate that *rgmb* mutants provide a sign stimulus exceeding that of wild-type, or that females are attracted to novelty, independent of specific color^2,45^. Consistent with the former possibility, females preferred wild-type over red-deficient *bdh1a-/-*, and even more strongly preferred *rgmb-/-* over *bdh1a-/-*.

Taken together, these analyses support a model in which redder coloration—imparted by carotenoids and preferred by females when choosing shoal mates—is associated with allelic variation across chromosomal regions that harbor BMP pathway and other genes expressed differentially between erythrophores and xanthophores, or across fin regions were these cells develop. During ontogeny of this trait (**Figure 4D**), progenitor cells experience different levels of BMP signaling that promote specification and differentiation as erythrophores (low levels) or xanthophores (intermediate to high) or that permit neither fate (highest). Because ligand gene expression is biased distally in the outgrowing fin, and developing chromatophores are equipped with several BMP receptors (including Rgmb), thresholds for fate lead to proximal–distal pattern. If signals are attenuated, locally produced cell fates are altered and pattern features are displaced. Importantly these events unfold in the context of growth and changes in positions^14^: some chromatophores are carried distally as the region of highest signaling grows distally; other chromatophores remain proximally as the fin tip grows away from them. These dynamics make it likely that developing chromatophores interpret thresholds for specification by integrating both signal amplitude and duration. Nevertheless we cannot exclude the possibility of cell-type specific activities of particular ligands or receptors that would uniquely specify one fate over another; nor in the context of our model can we rule out the possibility that females might prefer some other attribute of *rgmb* mutants over those of wild-type fish.

The developmental genetic dissection we have undertaken provides insights into the origins of a trait that can serve as a behavioral signal to conspecifics. Our demonstration that the pathway functions in multiple species to influence pattern further suggests a generalized mechanism. With this backdrop, one can envisage how historical contingencies across phylogenetic lineages have led to species-specific patterns, reflecting complements of available chromatophores, variation that has arisen for signal availability and chromatophore-lineage responsiveness, and the broader behavioral and ecological context of selection. Beyond fins and pigmentation, contributions of BMP signaling are evident for beetle horns and have been suggested for the comb of chickens^46^, antlers of deer^47^ and other exaggerated structures relevant to animal communication often occurring at anatomical extremities^48^. The pathway might be especially “co-optable” for such roles, given pre-existing functions in appendage outgrowth^22,49^ as well as skin appendage patterning (feathers, hair, scales)^50^, and the potential of this pathway for evolutionary fine-tuning of gradients, cell-type specific thresholds and allometric relationships among traits.

## Acknowledgements

Supported by NIH R35 GM122471 to DMP. Thanks to Parichy lab members for assistance with fish care, TA Larson and LB Patterson for helpful discussion and advice and Sarah Wilmsen and the University of Virginia Molecular Electron Microscopy core for assistance with electron microscopt.

## Author contributions

DMP and DH conceptualized the experiments. DH, TL, EHK, Y Liang and Y Liu conducted the experiments, supervised by DMP and JCC. JL contributed unpublished genomic annotations. DMP, DH, PLHdM, Y Lu and JCC analyzed the data. DMP and DH wrote the manuscript.

---

**Figure S1.**
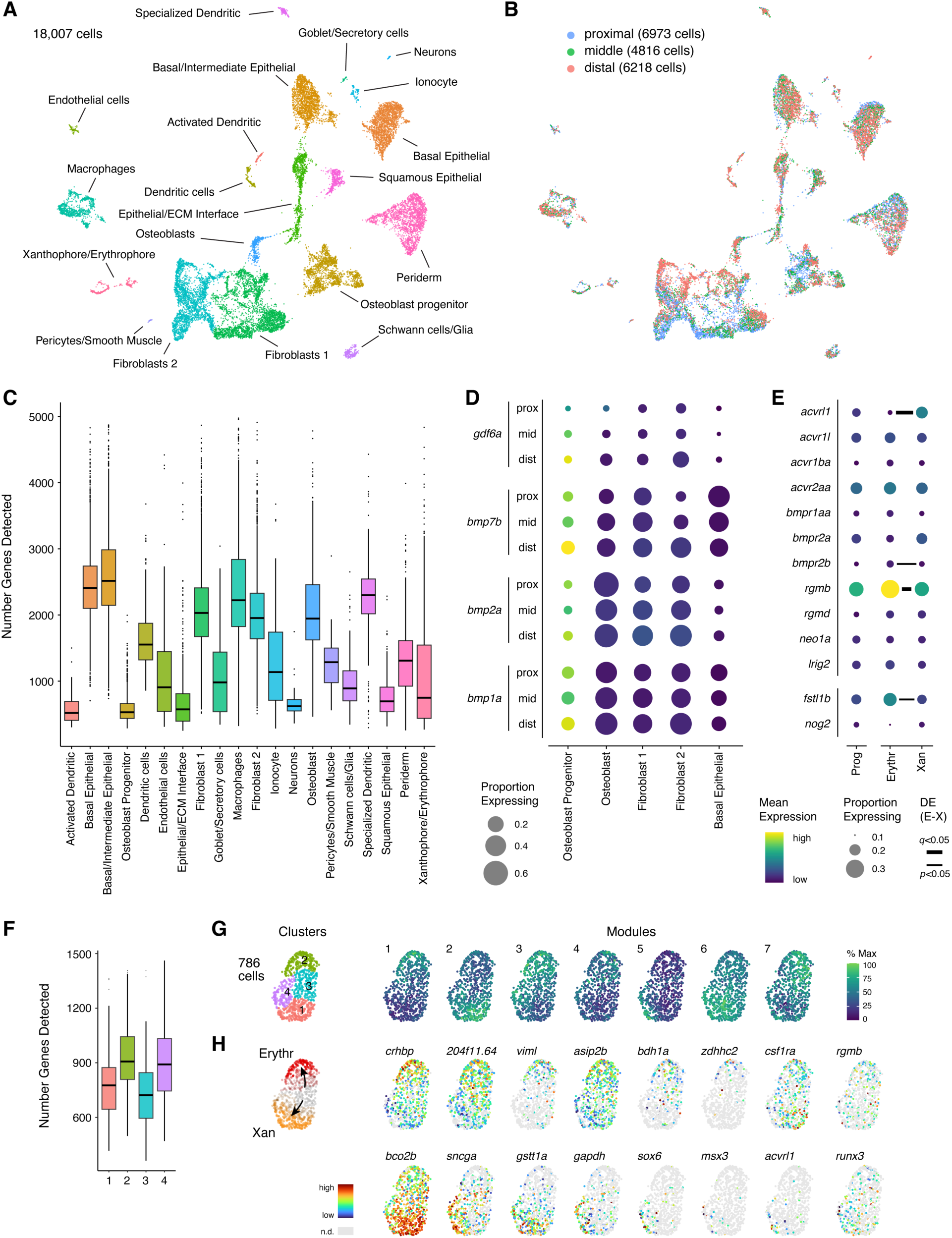
Transcriptomic analyses reveal differential expression of BMP ligand genes in fin tissue environment and modules of gene expression associated with xanthophore and erythrophore differentiation. (A–D) Results for cells isolated from proximal, middle and distal regions of wild-type anal fins at ∼12 mm standard length (SL) for 10X Chromium single cell RNA-sequencing (scRNA-Seq), subsetted for resident cells rather than circulating erythroid and immune cells (18,007 of 24,358 cell total). (A, B) Cluster cell type assignments and relative contributions of proximodistal cell origins, respectively. (C) Numbers of genes detected were variable across cell types, possibly reflecting different efficiencies of transcript recovery due to fragility of some cell morphologies (D) BMP ligand genes were differentially expressed proximodistally and also across cell types in the local tissue environment of pigment cells (D). Examples here had distal-biased expression overall (from Figure 1E), with especially high expression in dermal fibroblasts rich in extracellular matrix transcripts and likely to interact with pigment cells^1–3^. These data do not indicate if sparsity of expression across these cells reflects technical “drop-out,” due to lesser sampling than other cell-types, or variation due to cell-type specific biological processes (e.g., transcriptional bursting or other heterogeneities in cell subtypes or states). (E–H) Results for pigment cells and progenitors isolated by fluorescence activated sorting of *aox5*:nucEos-expressing^4^ cells from anal fins of 8–14 mm SL larvae for 10X Chromium scRNA-Seq. (E) Transcripts for canonical and non-canonical BMP receptors as well as secreted inhibitors of BMP signaling were evident amongst *aox5*+ cells representing unpigmented progenitors, erythrophores and xanthophores. Expression levels were assessed between erythrophores and xanthophores with significance levels of differences indicated by horizontal bars. (F) Numbers of genes detected among clusters of *aox5*+ cells. (G) Four clusters of *aox5*+ cells were detected with expression levels shown for each of 7 modules of co-expressed genes in UMAP space. (H) Inspection of cluster-specific and module-specific gene expression (Tables S3, S4), and comparison with results of bulk mRNA-Seq for mature cells (Table S1) indicated that clusters 1 and 2 represented xanthophores and erythrophores respectively, whereas clusters 3 and 4 were more likely to represent orange-pigmented^4^ or unpigmented cells not yet committed to either sublineage. Examples of some informative genes are mapped onto UMAP space.

**Figure S2.**
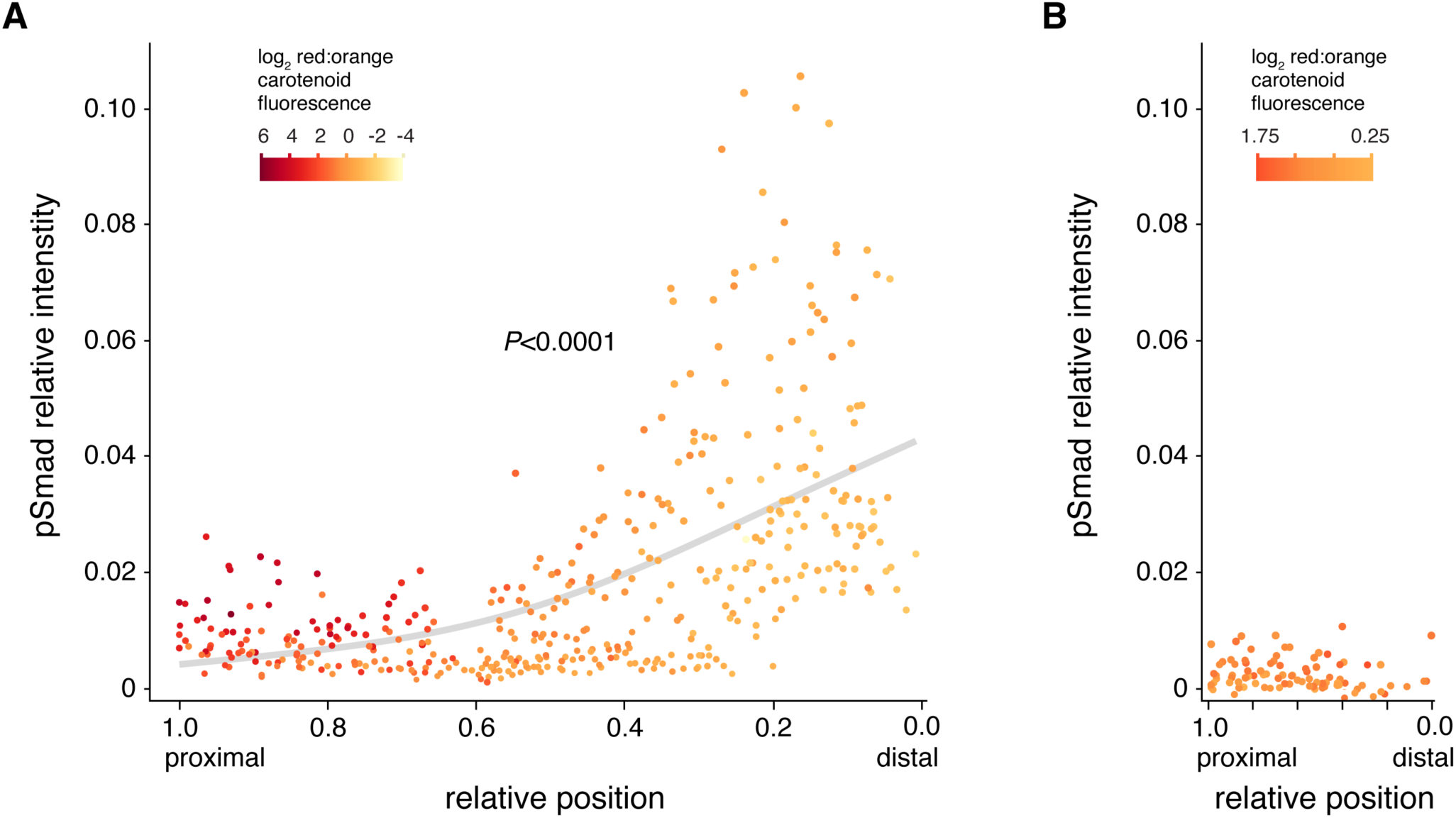
Graded BMP signal reception across the fin revealed by pSmad immunoreactivity. (A) Nuclear pSmad immunoreactivity relative to Dapi fluorescence in ∼15 mm SL fish when erythrophores and xanthophores have differentiated with differences evident in red-to-orange carotenoid fluorescence ratios (*n*=3 wild-type fish, 421 cells). pSmad immunoreactivity was significantly greater distally, among xanthophores, than proximally where erythrophores predominated (curve fit slope = −0.11, quadratic = 0.07; both parameters *P*<0.001). Positions are relative to most proximal and distal cells assessed. (B) At ∼8.5 mm SL only progenitors to erythrophores and xanthophores of intermediate color are present^4^ and no gradient in pSmad immunoreactivity was evident (quadratic and linear models, both *P*>0.15; *n*=3 fish, 90 cells). Plot widths in A and B are proportional to proximal–distal fin lengths at the two stages shown.

**Figure S3.**
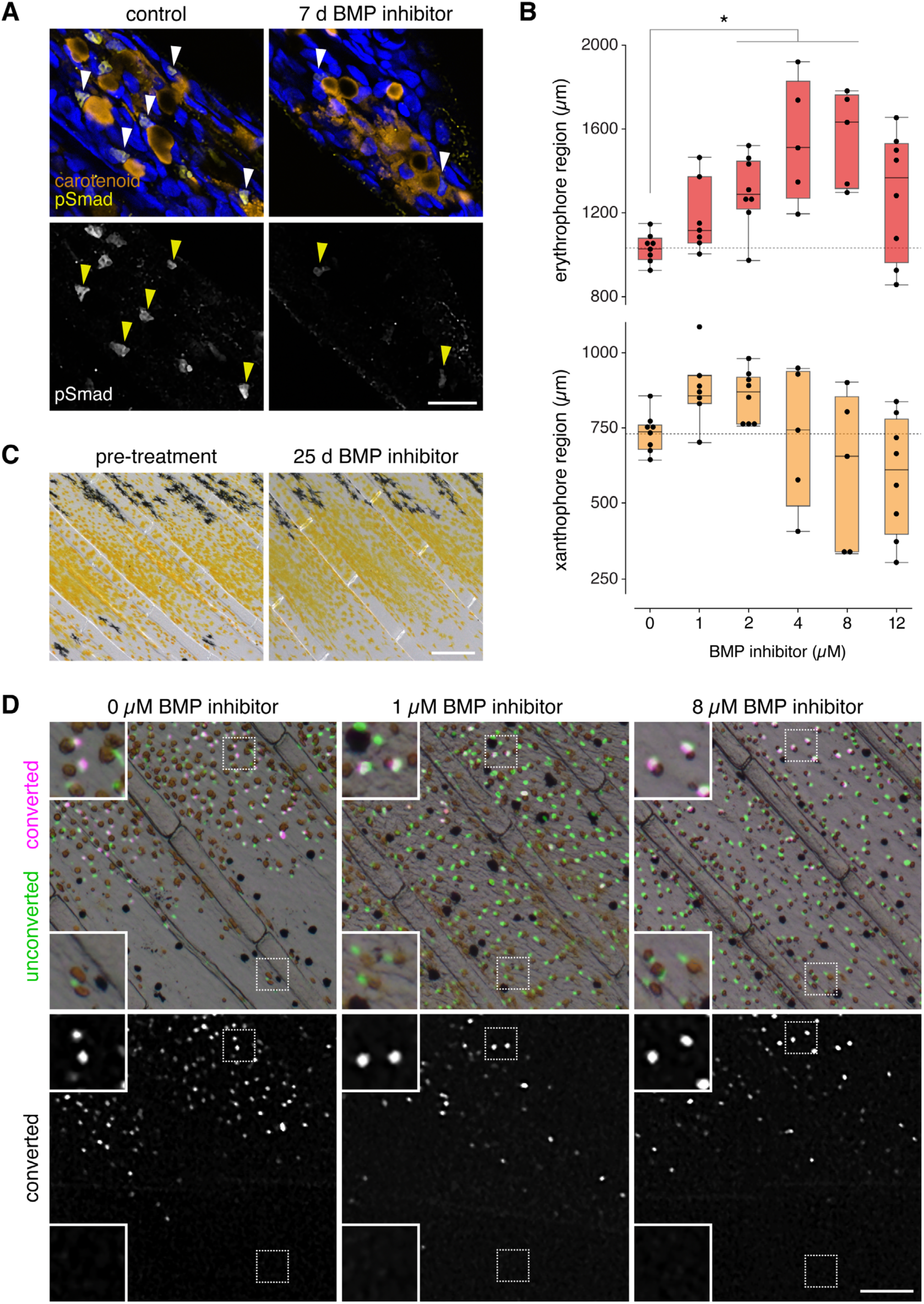
BMP signal attenuation affected differentiation. (A) pSmad immunoreactivity was pronounced in nuclei of xanthophores at the distal portion of the fin, yet BMP inhibitor markedly attenuated this staining. Arrowheads show examples of nuclei adjacent to carotenoid accumulations with either strong immunostaining (control) or weak immunostaining (7d BMP inhibitor). Orange, xanthophore carotenoid autofluorescence; yellow, pSmad staining; blue, Dapi. (B) Regions covered by erythrophores and xanthophores were altered by BMP inhibitor treatment, as measured by proximal–distal distances populated by these cells at the fifth fin ray. BMP inhibition led to significantly larger regions covered by erythrophores (treated vs. control, contrast *F*_1,35_=16.8, *P*=0.0002; *post-hoc* means comparisons to control: *, Dunnet’s test *P*<0.05; overall ANOVA *F*_5,35_=5.2, *P*=0.0011). Regions covered by xanthophores varied with treatment (overall ANOVA *F*_5,35_=3.9, *P*=0.0066) but could be greater than controls, particularly at low inhibitor doses, or less than controls at high doses. Inhibitor treatment also led to significantly increased variability in coverage by erythrophores and xanthophores (Levene’s tests *P*=0.0073, *P*=0.0009, respectively), leading to heteroscedasticity in residuals, adequately corrected for parametric analyses by reciprocal transformation (erythrophore regions) and square transformation (xanthophore regions). (C) Xanthophores that had already developed in older fish were similarly abundant before treatment and after 25 d of BMP inhibitor treatment, suggesting xanthophores do not die as a consequence of BMP signaling inhibition. (D) Fate mapping with photoconvertible fluorophore expressed by xanthophores and erythrophores (*aox5*:nucEosFP) showed that cells marked by photoconversion (green ➝ red, shown here in magenta for visual accessibility) early in pattern development and fin outgrowth did not migrate to distal regions where control fish have very few carotenoid-containing pigment cells (0 *µ*M), fish with low doses of inhibitor developed xanthophores (1 *µ*M) and fish with high doses (8 *µ*M) developed erythrophores. Presence of only unconverted green fluorophore in distal regions further indicated that cells had not yet started to express *aox5*:EosFP and so had differentiated from unspecified progenitors, likewise excluding the possibility of transdifferentiation, which has been documented unidirectionally (erythrophores ➝ xanthophores) during regeneration^4^. Upper panels show brightfield images superimposed on fluorescence of unconverted fluorophore and converted fluorophore. Continued expression of *aox5*:nucEosFP after conversion leads to all cells having unconverted fluorophore and appearing white (green+magenta) except where live specimen movement during acquisition led to channel offset. Scale bars, 20 *µ*m (A), 200 *µ*m (C), 100 *µ*m (D).

**Figure S4.**
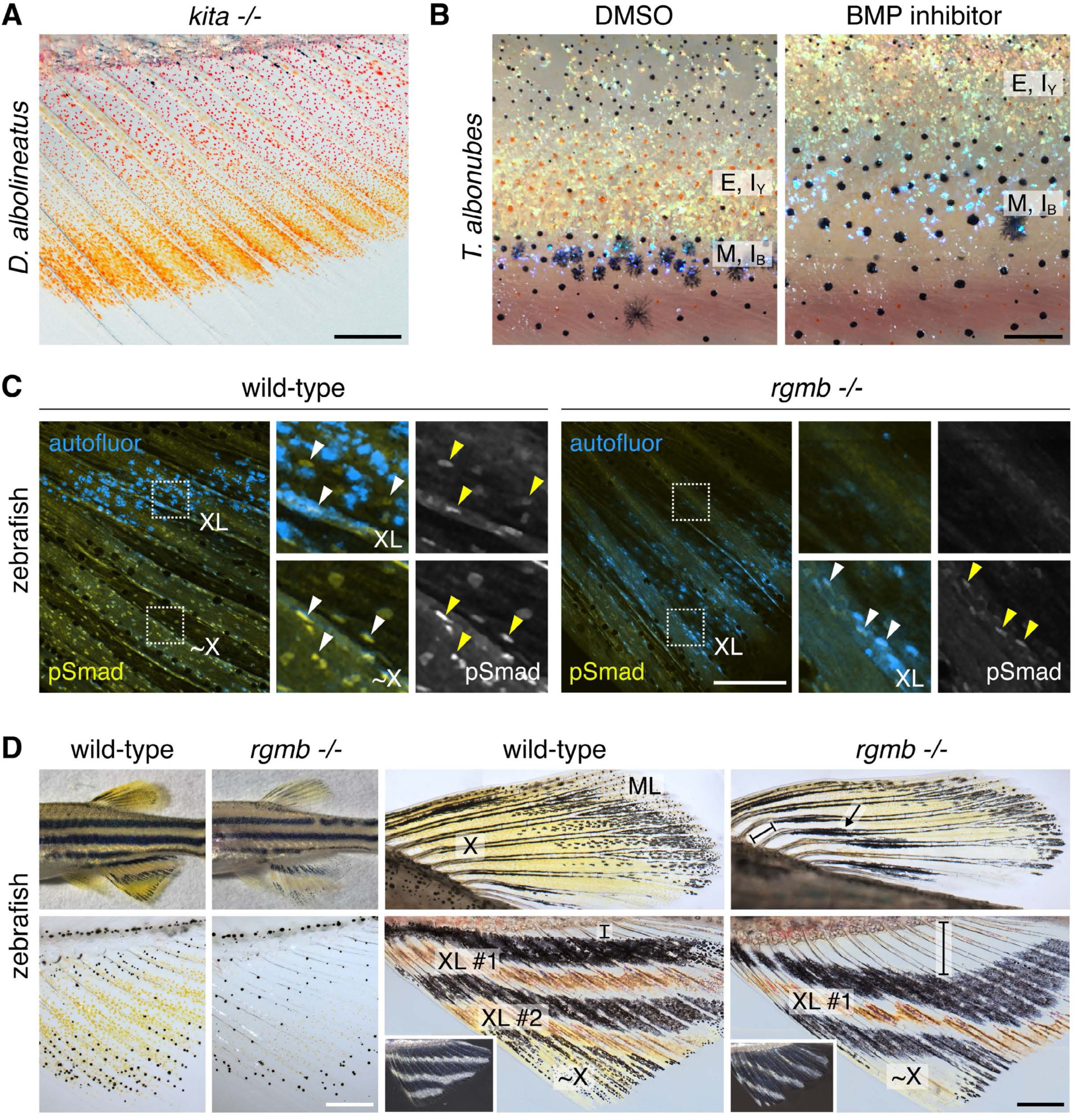
Additional roles for BMP signals in setting pattern phenotype. (A) In *D. albolineatus*, neither the proximal boundary between erythrophores and xanthophores nor the distal boundary of xanthophores depended on melanophores normally at these locations, as shown here in a mutant for the Kit receptor tyrosine kinase encoded by *kita*, which lacks fin melanophores due to an autonomous requirement for Kit in this lineage^5,6^. Compare with Fig. 1A. (B) A BMP requirement for body pigment pattern of *T. albonubes* was revealed by inhibitor-treated fish, in which regions of erythrophores and yellow iridophores (E, I_Y_) and melanophores and blue iridophores (M, I_B_) were shifted dorsally in comparison to controls. Yellow and blue iridophore morphologies and arrangements resembled those of zebrafish interstripes and stripes, respectively.^7^ (C) In wild-type zebrafish, xantholeucophores (XL) were only weakly pSmad+ (autofluorescing pigmentary contents adjacent to pSmad+ nuclei), whereas more distal xanthophore-like cells (∼X) were strongly pSmad+. In the *rgmb* mutant, xantholeucophores were shifted distally and pSmad immunoreactivity was reduced overall. (D) In comparison to the wild type, *rgmb* mutants had moderately more frequent stripe breaks and reduced yellow pigmentation (upper left; ∼13.5 mm SL) and xanthophores in the fins were markedly delayed in their differentiation (lower left; ∼9 mm SL). In dorsal fins of young adults (∼3 months old), a pigment-cell free region (bracket) and ectopic melanophore stripe were present where xanthophores (X) occurred in wild-type; melanoleucophores (ML) were markedly fewer^3^. In anal fins, the first developing interstripe of xantholeucophores (XL #1) was present distal to the corresponding interstripe of wild-type and a larger pigment-cell free region was evident proximally (brackets). In older *rgmb* mutants additional interstripes eventually formed, with angled or irregular patterns (insets, ∼8 months old). Scale bars, 500 *µ*m (A), 200 *µ*m (B,C), 2 mm (D upper left), 250 *µ*m (D lower left), 500 *µ*m (D right).

**Figure S5.**
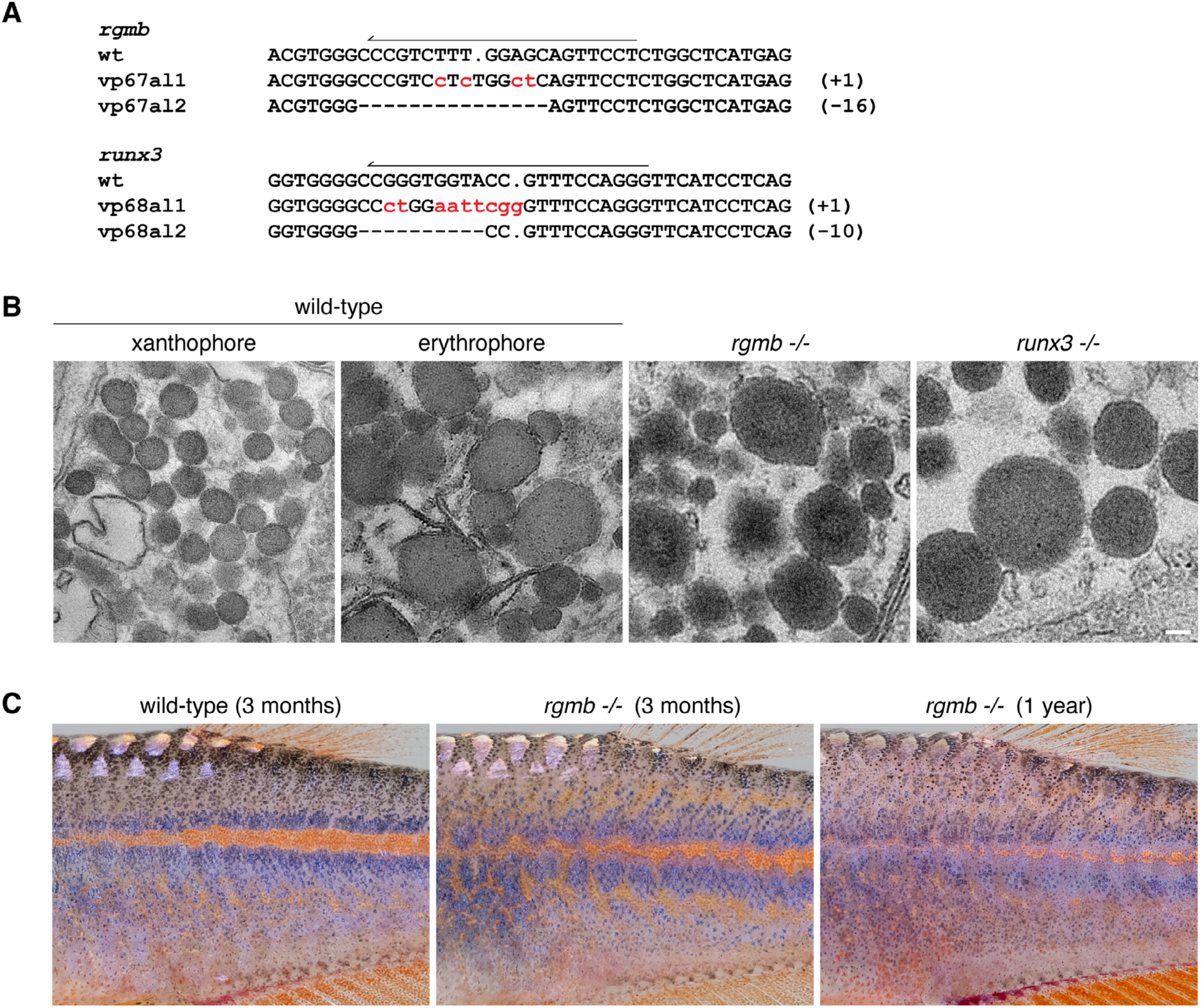
Mutants for *rgmb* and *runx3* of *D. albolineatus*. (A) Frameshift alleles recovered following CRISPR/Cas9 targeting that that led to premature termination codonsl differences in phenotype were not evident across alleles. Arrows indicate CRISPR target sites in wild-type with numbers at right showing total numbers of base pairs inserted or deleted. (B) Transmission electron microscopy showing electron-dense carotenoid vesicles of fin distal xanthophores and proximal erythrophores in wild-type as well as ectopic distal erythrophores in *rgmb* and *runx3* mutants. Vesicles of mutant cells were of similar size to those of erythrophores in wild-type, which were consistently larger than those of xanthophores. (C) Body pigmentation of rgmb mutants differed from wild-type in having a somewhat less distinctive and more irregular pattern of already weak melanophore stripes and light interstripe. During later development the pattern of wild-type was maintained whereas an increasingly diffuse pattern emerged in *rgmb* mutants. Image of 1-year fish is re-scaled to 75% of original size to illustrate an anatomical region comparable to that of 3-month fish). Scale bar, (B) 100 nm.

**Figure S6.**
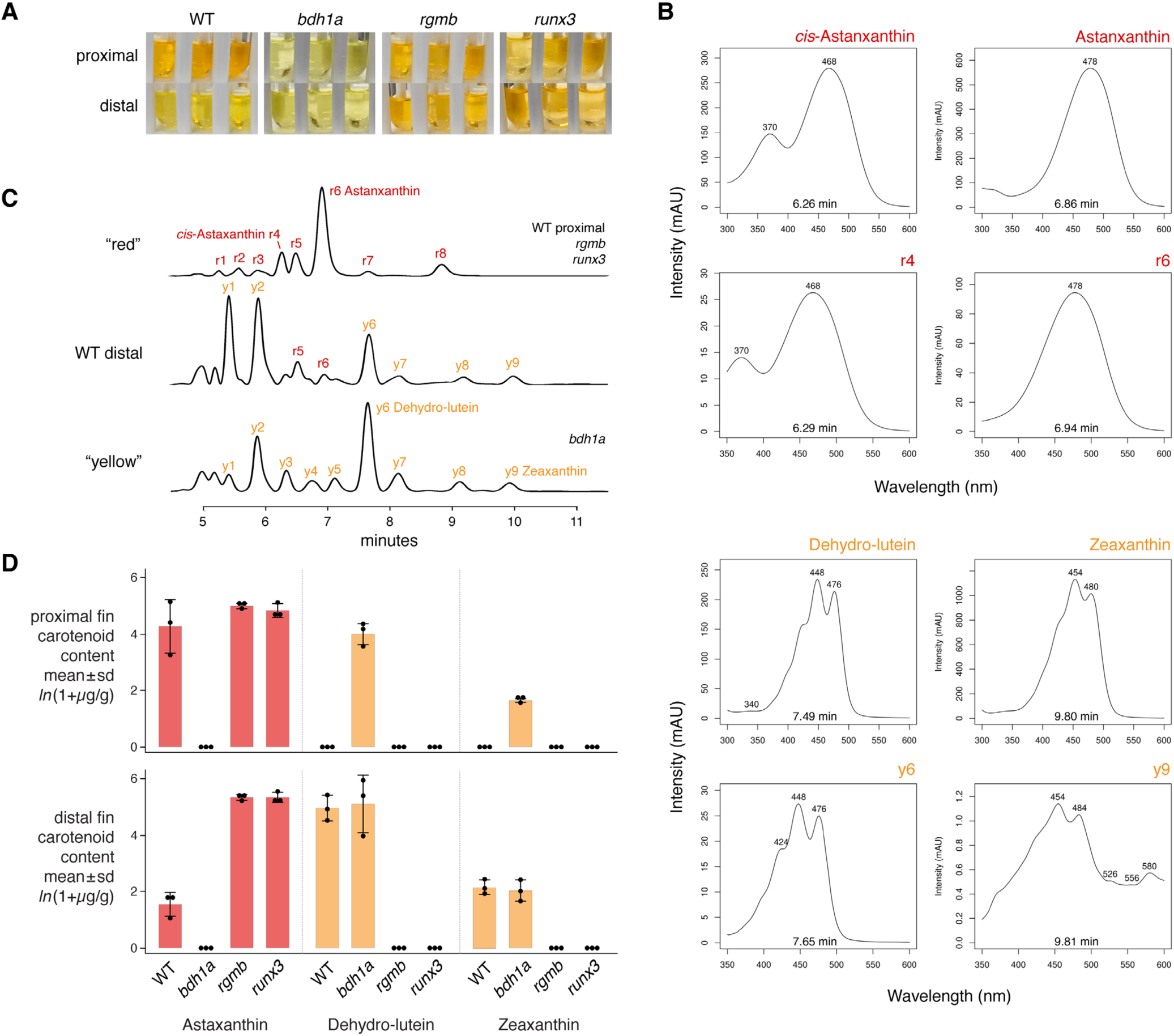
Chemical analysis of carotenoid contents. (A) Color differences were evident during extraction of carotenoids from proximal and distal fin regions of wild-type but not mutants. Each anatomical pair of samples represents fin tissue pooled from 6 individual fish. (B) Standards for carotenoids (red: *cis*-Astaxanthan, Astaxanthan; yellow: Dehydro-lutein, Zeaxanthin) above inferred corresponding peaks from red or yellow tissue in C. (C) Example chromatograms of exclusively red or yellow tissue (representative of genotypes and region shown at right), along with distal tissue of wild-type with inferred unique and overlapping peaks numbered. (D) Quantification of tissue contents for identified carotenoids (ANOVA genotype x position interactions all *F*_3,16_>23, *P*<0.0001 after *ln*-transformation).

**Figure S7.**
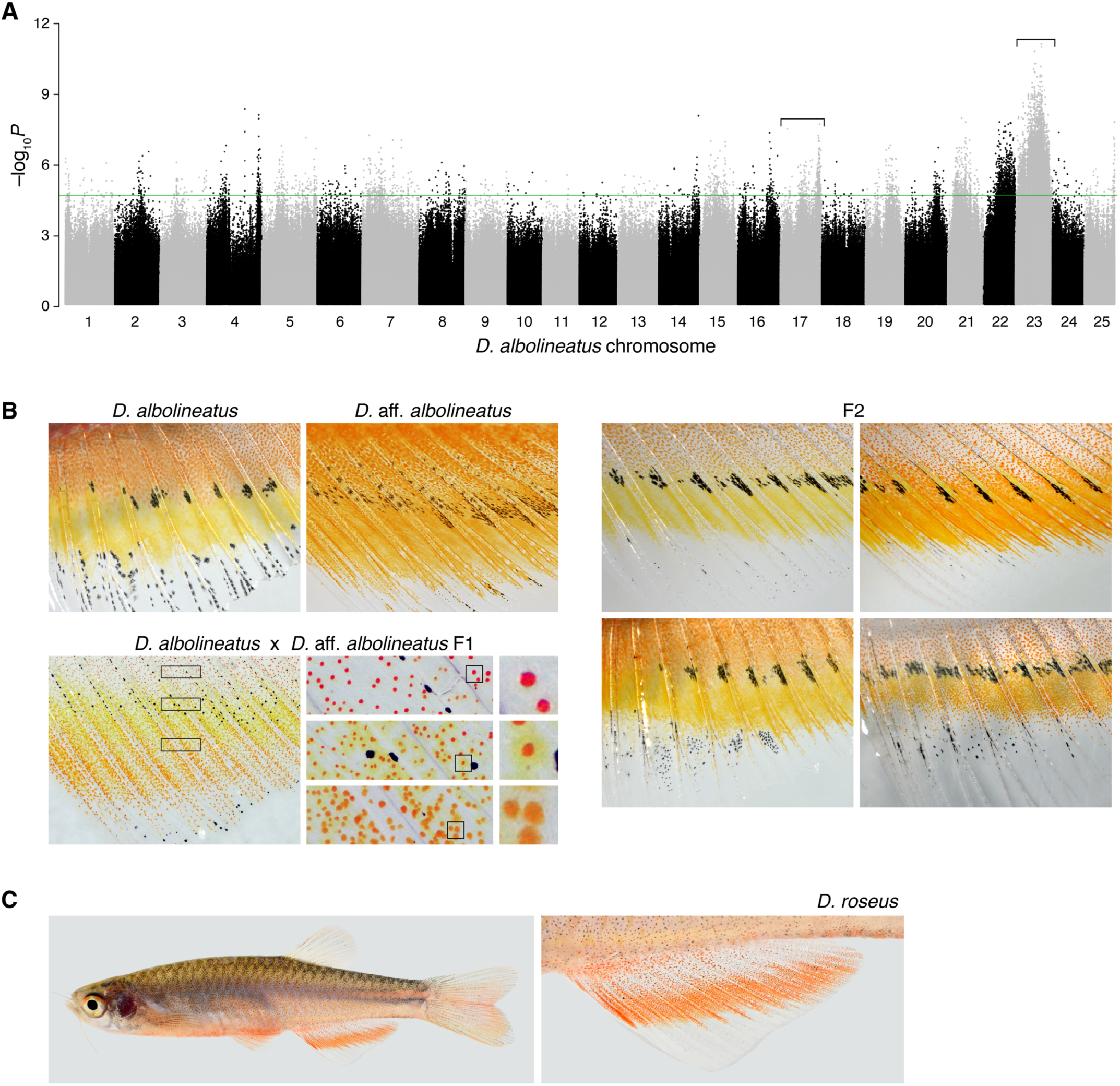
Fin pattern variation in *D*. aff. *albolineatus*, *D. albolineatus* and related taxa. (A) A naturally occurring variant in fin pigmentary phenotype was associated with alleles on Chr23 and other chromosomes. Manhattan plot shows Fst values across genome (*P*-values Fisher’s Exact test; green threshold, top 0.1%). Details of Chr17 and 23 shown in Figure 4B. (B) Phenotypes of fish used in genetic mapping, showing males of parental strains *D. albolineatus* and *D.* aff. *albolineatus*, along with fin pattern of the F1 male, and examples of F2 progeny. In the F1, chromatophores especially in the middle of the fin had prominent concentrations of red carotenoids following epinephrine treatment and also a widely dispersed yellow pigment. *D. albolineatus* xanthophores and erythrophores have pteridines that are colorless to the human eye^4^ and it is conceivable that a yellow pteridine is expressed in *D.* aff. *albolineatus* or uniquely in the hybrid. (C) Origins and phlogenetic history of alleles within *D.* aff. *albolineatus* are uncertain. Nevertheless, widespread hybridization and allelic introgression elsewhere in the genus^8^ raise the possibility that alleles for fin color and pattern in *D.* aff. *albolineatus* might derive from a closely related species of subspecies. In this context *D. roseus* is notable as this species is superficially similar to *D. albolineatus*, yet lacks prominent yellow fin coloration as in *D*. aff. *albolineatus*. *D. albolineatus* and *D. roseus* are sympatric with both found over considerable geographical areas that include Myanmar, Thailand, Laos and the island of Sumatra, Indonesia^9–11^.

**Figure S8.**
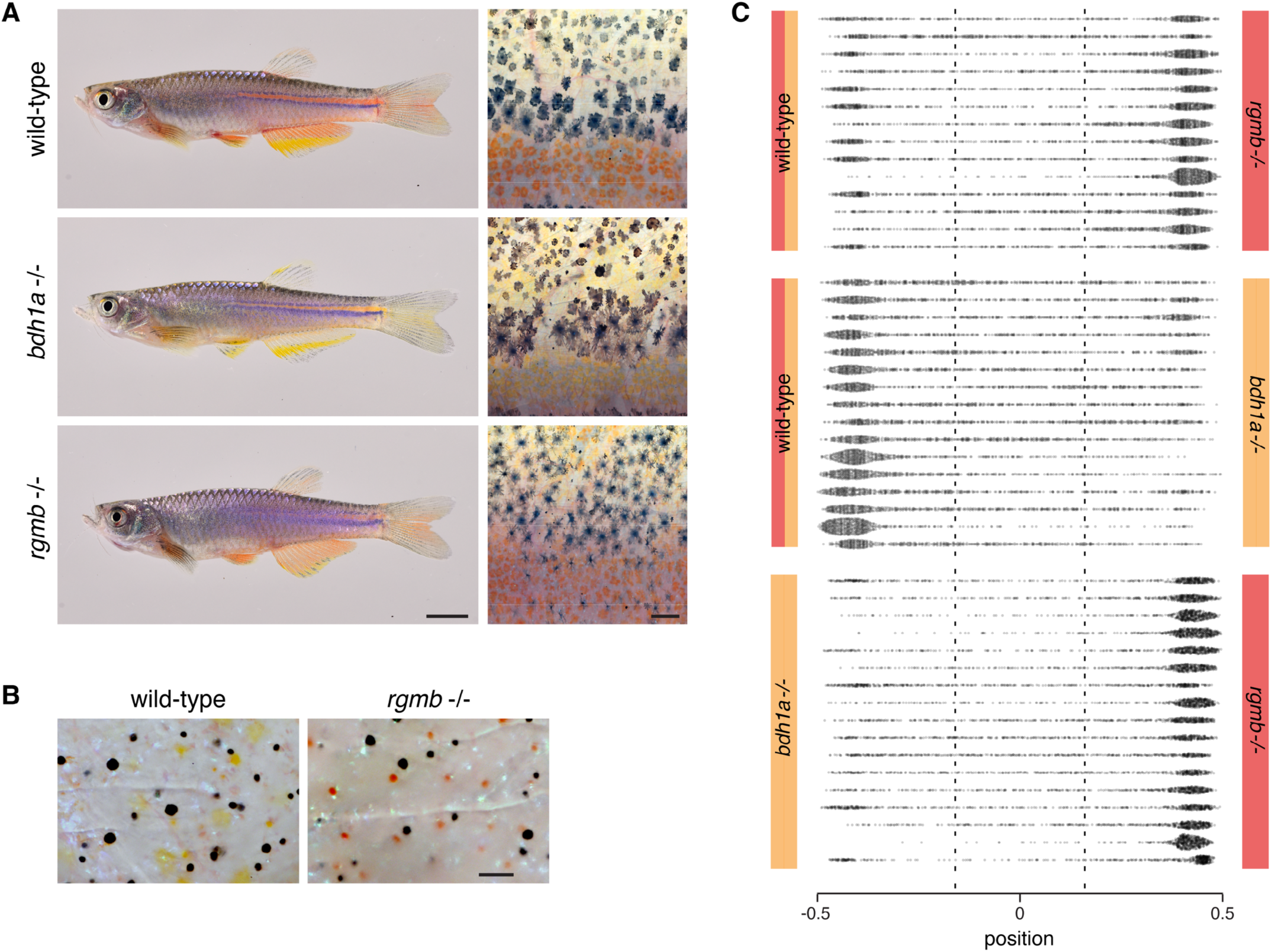
Red-deficient and super-red phenotypes provide opposing signals for shoaling. (A) Whole-fish phenotypes and flank closeups of wild-type, *bdh1a* and *rgmb* mutants. Higher magnification views at right illustrate patterns and color differences in the vicinity of a residual dorsal melanophore stripe in wild-type (darker melanophores). Ventral to this stripe are cells of the interstripe, comprising erythrophores and deeper iridophores that appear as a blue-tinted matte of cells in this image. Dorsal to the stripe are lighter melanophores interspersed with xanthophores. In *bdh1a-/-*, yellow cells occur both within the interstripe and dorsally. In *rgmb-/-*, erythrophores of the interstripe are more numerous, melanophores are more dispersed and lighter in color, and reddish erythrophores are found in place of xanthophores dorsally. (B) Body pigment cells at dorsal scales of wild-type and *rgmb* mutant following epinephrine treatment, showing differences in pigment color. (C) Positions of individul female fish relative to stimulus males as determined at 0.5 sec (15 video fame) intervals. Each row is a different individual; actual sides used for stimulus shoals were alternated betweem trials. Vertical dashed lines demarcate thirds of each tank with preference areas defined for each stimulus shoal at left and right (differences in total times spent between areas are shown in Figure 4C). Scale bars, 5 mm (A, left), 200 *µ*m (A, right), 100 *µ*m (B).

## STAR★METHODS

### KEY RESOURCES TABLE

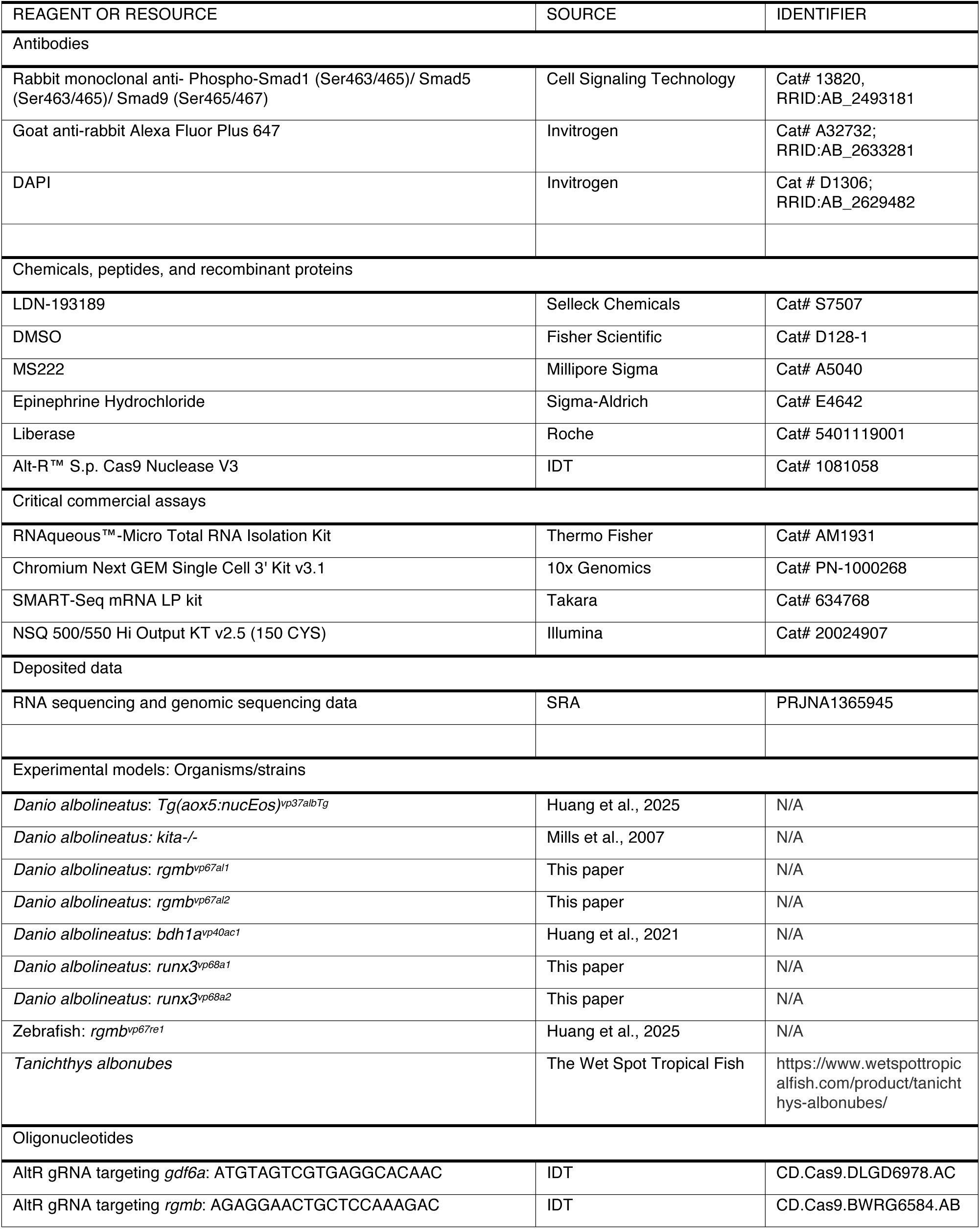
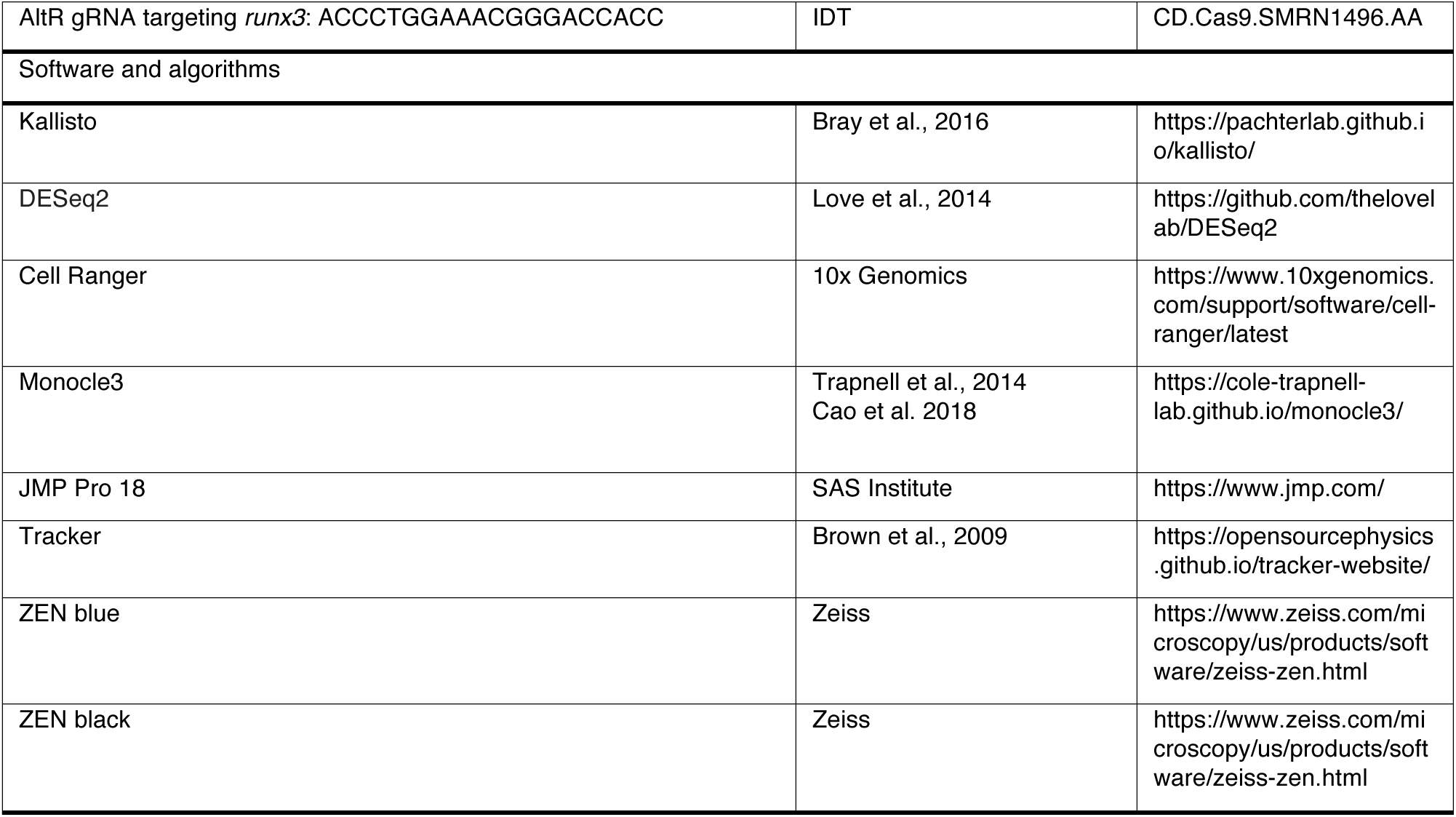

### EXPERIMENTAL MODEL AND STUDY PARTICIPANT DETAILS

*Danio albolineatus* were derived from individuals collected in Thailand by M. McClure in 1995, subsequently provided to the laboratory of S. Johnson (dec.), and have been maintained as a closed stock in our facility since 2000. *D*. aff. *albolineatus* were obtained from the pet trade in 2003 and have been maintained as a closed stock in our facility since that time. *Tanichthys albonubes* were obtained from The Wet Spot Tropical Fish (Portland, USA). Fish were reared under standard conditions used to maintain *D. rerio* (∼28 °C; 14 L:10D) with larvae initially fed marine rotifers derived from high-density cultures and enriched with Rotimac and Algamac (Reed Mariculture), with older larvae and adults subsequently fed live brine shrimp and a blend of flake foods enriched with dried spirulina, Tetramin and Z-Pro 350 (Zeigler Bros., Inc., USA). Stocks of mutant or transgenic fish included *D. albolineatus*, *rgmb^vp67al1^*, *rgmb^vp67al2^*, *runx3^vp68a1^*, *runx3^vp68a2^*, *bdh1^vp40ac1^* and Tg(*aox5:nucEosFP*)*^vp43aTg^*; *D. rerio rgmb^vp67re1^*. Developmental staging followed ref. ^17^, with minor modifications for use in *D. albolineatus*. Unless otherwise specified, male fish were used owing to the sexual dimorphism of *D. albolineatus*^4^. This study was performed in strict accordance with the recommendations in the Guide for the Care and Use of Laboratory Animals of the National Institutes of Health. All animals were handled according to approved institutional Animal Care and Use Committee (ACUC) protocol (#4170) of the University of Virginia. Euthanasia was accomplished by overdose of MS222 followed by physical maceration.

### METHOD DETAILS

#### CRISPR/Cas9 mutagenesis

CRISPR/Cas9 mutants or F0 mosaics were generated by injecting one-cell stage embryos with approximately 1 nanoliter of 5 μM gRNA:Cas9 RNP complex (IDT, AltR CRISPR/Cas9 reagents), which allowed for highly efficient mutagenesis^18^ even in F0 fish. For F0 mosaic analysis, 8 fish with slowed growth or eye defects (observed in the zebrafish mutant *gdf6a^vp65re1^*) were selected from 46 survivors and genotyped to assess CRISPR/Cas9 mutagenesis efficiency. To produce mutant lines, individual fish were sorted for anal fin phenotypes at juvenile stage and alleles were recovered by intercrossing and outcrossing.

#### Pharmacological Analysis

LDN-193189 (S2618, Selleck Chem) was used to inhibit BMP signaling. LDN-193189 was dissolved in DMSO (final DMSO concentration 0.5%). In Figure 2B–D, *D. albolineatus* were treated beginning at 8.5 mm standard length (SL) for a duration of 3 weeks. The concentration of LDN-193189 was set in a gradient of 0, 1 μM, 2 μM, 4 μM, and 12 μM. Control fish received 0.5% DMSO only. In Figure 3B, 8 mm SL *T. albonubes* were treated with either 4 μM LDN-193189 in 0.5% DMSO or 0.5% DMSO alone for 2 weeks. In both experiments, fish were transferred daily into fresh water during the day and fed live brine shrimp, then immersed in freshly prepared drug solutions overnight. For *D. albolineatus*, each treatment group contained 8 fish, except for the 12 μM condition which had 16 fish divided into 2 groups. By the end of the experiment, 1 fish in the 1 μM group, 3 fish in the 4 μM and 8 μM groups, and 8 fish in the 12 μM group died. Brightfield and fluorescence images were collected for the *D. albolineatus* experiment; 1 fish from the 2 μM group was excluded due to sample drift during fluorescence imaging. In the *T. albonubes* experiment, both groups initially contained 9 fish, and 2 fish died in the drug-treated group. For the chronic treatment assay in adult *D. albolineatus*, 4 fish were exposed to 4 μM LDN-193189 in 0.5% DMSO or to 0.5% DMSO alone for 7 days to assess drug efficacy. An additional 12 fish were imaged before and after undergoing the same drug treatment for 25 days. Fish were transferred to fresh water every 3 days to receive a single feeding of live brine shrimp before being returned to newly prepared drug solutions. No mortality occurred during the treatment.

#### Carotenoid analyses

Distal (xanthophore-containing) portions of the anal fins from adult *D. albolineatus*, *rgmb*, or *runx3* mutants were dissected and pooled for pigment extraction. 3 biological replicates were prepared, each consisting of 6 fin fragments. 200 *µ*l MilliQ water, 300 *µ*l 200-proof ethanol and 600 *µ*l hexane:tert-butyl methyl ether (hexane:MTBE, 1:1, vol:vol) were added to fish fin samples and mixed by vigorous vortexing. Samples were centrifuged at 12000 g for 5 min, and the supernatants were collected. Extraction was repeated until the fins turned pale. The combined supernatants were dried under nitrogen gas, then saponified with 0.02 M NaOH in ethanol for 4 h. 1 ml saturated NaCl, 2 ml MilliQ water and 2 ml MTBE were added to the saponified extract, then the samples were vortexed and centrifuged at 2000 g in a glass tube. Supernatants were collected and dried under nitrogen gas, before being redissolved in the mobile phase for HPLC analysis. The extracted fish fins were dried and weighed to obtain dry weight measurements. To perform reverse-phase HPLC analysis, carotenoids were dissolved in acetonitrile : methanol : tetrahydrofuran (48:48:4, vol:vol:vol) and injected into an Agilent 1200 series HPLC fitted with a YMC carotenoid column (5.0 μm, 4.6 mm × 250 mm). Samples were then eluted with a gradient mobile phase of acetonitrile:methanol:tetra-hydrofuran (45:45:10, vol:vol:vol) through 9 min, ramping up to acetonitrile:methanol:tetra-hydrofuran (35:35:30) from 9–10 min, followed by isocratic conditions through 12 min, before ramping back down from 12-13 min, followed by isocratic conditions through 15 min. Solvent was pumped at a constant rate of 1.2 mL/min, and the column was maintained at 30°C. Sample elution was monitored using a UV-Vis photodiode array detector at various wavelengths. Carotenoid peaks were identified by comparison to authentic standards of astaxanthin, zeaxanthin, lutein, and dehydro-lutein, as well as their putative *cis* isoforms. Consistent peaks were quantified using three methods: peak area of total signal between 350-600 nm normalized to dry weight; peak area of total signal between 350-600 nm normalized to the combined signal of all peaks; and by absolute quantification using curves generated from authentic standards. We also used fluorescence imaging to measure red vs. yellow carotenoid ratios as described^4^. Briefly, laser intensities for red (excitation wavelength 561 nm) and green (excitation wavelength 488 nm) channels were set to be identical. The ratio of red-to-green signal was quantified to obtain an estimate of the relative of abundance of red vs. yellow carotenoids. For BMP inhibitor experiments, the most distal pigmented cells in the center of the anal fin were imaged. For pSmad immunoreactivity experiments, cells along the 4^th^ fin ray were imaged.

#### Immunohistochemical staining

Fins were fixed in 4% paraformaldehyde (PFA) for 1 hour at room temperature and washed 5 times for 5 minutes each with PBST (0.1% Triton X-100 in 1× PBS). Samples were then incubated overnight in blocking solution (2% goat serum in PBST), followed by incubation with a primary antibody against phosphorylated Smad1/5/9 (pSmad; 1:200 dilution, #13820S, Cell Signaling Technology) overnight at 4 °C. After washing 5 times with PBST for 5 minutes each, samples were incubated overnight with a fluorescent dye–conjugated secondary antibody (goat anti-rabbit Alexa Fluor Plus 647, 1:400), washed again, and imaged. Three adult *D. albolineatus* and *T. albonubes* were analyzed. Among the 14 offspring of the *rgmb^vp67re1^*^/+^ x *rgmb^vp67re1/vp67re1^* cross, 8 heterozygotes and 6 homozygotes were analyzed.

#### Lineage tracing analysis

Photoconversions were performed on *Tg(aox5:nucEos)^vp37albTg^* fish using a Zeiss LSM 800 laser-scanning confocal microscope equipped with a 405 nm laser and ZEN Blue software. A small rectangular region containing the most distal *nucEos*⁺ cells was photoconverted with 405 nm laser in the anal fin of 9 mm SL larvae. Then fish were treated with 0.5% DMSO, 1 μM LDN-193189 in 0.5% DMSO, or 8 μM LDN-193189 in 0.5% DMSO for 2 weeks (*N* = 8; no exclusions). Following photoconversion, fish were maintained in tanks shaded from ambient light to prevent spontaneous photoconversion. Final imaging session was performed with fluorescence and then brightfield mode.

#### Imaging and image processing

For the whole fish imaging, fish were euthanized and captured on a Nikon D-810 digital single lens reflex camera with MicroNikkor 105 mm macro lens. Anal fin details were imaged using a Zeiss Axio Observer inverted microscope or Zeiss AxioZoom stereomicroscope equipped with Zeiss Axiocam cameras. Carotenoid autofluorescence was imaged using a Zeiss LSM880 inverted laser confocal microscope in Airyscan SR mode. Images were acquired using a Zeiss AxioObserver inverted microscope equipped with Yokogawa CSU-X1 M5000 laser spinning disk and Hamatsu camera. Images were captured either as single frames or as tiled sets of larger areas that were then stitched computationally using ZEN Blue or ZEN Black software. Color balance and display levels were adjusted for entire images as needed and kept consistent across comparison. Signal intensity and nuclear distance from fin tip were measured using ZEN Black software.

#### Transmission Electron Microscopy (TEM)

Fins were amputated and pre-fixed for 15–30 min with 4% formaldehyde and 2.5% glutaraldehyde in PBS, then transferred to fresh 4% formaldehyde and 2.5% glutaraldehyde in 100 mM phosphate buffer (pH 7.4) for an additional 2 hr at room temperature. Tissues were post-fixed for 2 hr with 2% osmium tetroxide and 2% uranyl acetate at room temperature, dehydrated in acetone, and flat-embedded in epoxy resin. Embedded blocks were trimmed to create a pyramid shape for ultramicrotomy. Seventy-nanometer sections were cut, collected on carbon mesh grids, and post-stained with lead citrate and Uranyless™. Imaging was performed on an FEI Tecnai F20 equipped with a TEITZ XF416 detector.Bulk RNA-Seq and Single-cell RNA-Seq

#### Bulk RNA-Seq and Single-cell RNA-Seq

∼100 male *Danio albolineatus Tg(aox5:nucEosFP*)*^vp37albTg^* of 1.5 months old were euthanized. Anal fins were dissected and tissue collected from proximal erythrophore or distal xanthophore regions in PBS and dissociated to cell suspension by Liberase (Roche, Cat# 5401119001) treating at 30 °C for 30 min and filtering through 40 μm strainer. Isolated cells were sorted through Sony SH800 fluorescence activated cell sorter (Sony, Japan). GFP+ healthy cells were collected, and RNA was extracted using RNAqueous™-Micro Total RNA Isolation Kit (Thermo Fisher, Cat# AM1931). Sequencing libraries were constructed using SMART-Seq mRNA LP kit (Takara, Cat# 634768) and sequenced on an Illumina Nextseq-550 using NSQ 500/550 Hi Output KT v2.5 (150 CYS). Reads were aligned to *Danio albolineatus* reference genome using Kallisto and analyzed using DESeq2. For single-cell RNA-seq, ∼150 anal fins were collected from *Tg(aox5:nucEosFP*)*^vp43aTg^* fish of 8 mm SL to 14 mm SL and sorted for GFP+ cells. Library production was performed using a Chromium controller (10X Genomics, USA) and Chromium Next GEM Single Cell 3’ Kit v3.1 (10X Genomics, USA). Quality control and quantification assays were performed using a Qubit fluorometer (Thermo Fisher, USA) and a 2100 Bioanalyzer (Agilent, USA). We built a zebrafish STAR genome index using Lawson Lab zebrafish transcriptome annotation(58), filtered for protein-coding genes. Final cellular barcodes and Unique Molecular Identifiers (UMIs) were determined using Cell Ranger (10X Genomics, USA). Data were analyzed in Monocle3. In brief, we filtered cells for quality to have less than 5% mitochondrial reads, and greater than 500 unique molecular identifiers and 200 genes expressed. RNA-seq data are available through SRA (accession ID: PRJNA1365945). We used Uniform Manifold Approximation and Projection (UMAP) to project transcriptomic space in two dimensions followed by Louvain clustering. We assigned clusters to cell types by comparing genes detected to published cell type specific markers. Cell type marker analysis, differential expression and trajectory analyses used standard methods and functions as described (cole-trapnelllab.github.io/monocle3/docs/). Gene module identification was performed using Monocle3 find_gene_modules() across clusters.

#### Genome Assembly and Annotation

For assessing standing genetic variation in fin phenotype we first generated a high quality PacBio genome assembly for *D. albolineatus*. To obtain a more contiguous assembly, we *de novo* assembled the *D. albolineatus* genome. High-fidelity (HiFi) reads were generated from a single individual of *Danio albolineatus* from the Parichy lab. Raw sequencing reads in BAM format were filtered to retain only those with a quality score (Read Quality, RQ) greater than Q20 (RQ >= 0.99) using bamtools. The filtered BAM files were then merged and converted to FASTQ format using pbindex and bam2fastq (PacBio SMRT Tools). Read quality was assessed using NanoPlot. The pre-processed HiFi reads were assembled *de novo* using Hifiasm (v0.19-r320) with default parameters. The resulting primary contig assembly (*.bp.p_ctg.gfa) was extracted and converted to FASTA format using awk. The initial *de novo* assembly was assessed for completeness using QUAST (v5.0.2) and BUSCO (v5.4.3) with the Actinopterygii lineage dataset. To improve the contiguity and order the contigs into chromosome-scale scaffolds, the assembly was polished and scaffolded using RagTag (v2.1.0). Given the conservation on karyotype number across *Danio*, the high-quality, chromosome-level assembly of the reference species, *Danio rerio* (NCBI GCF_000002035.6, GRCz11), was used as the reference guide for scaffolding. Repetitive elements were identified and masked in a multi-step procedure. First, repeats were identified *de novo* using RepeatModeler (v2.0.4) on the scaffolded genome to build a custom repeat library. The assembly was then masked in three iterative rounds using RepeatMasker (v4.1.2): 1. Simple Repeats: Masking for simple repeats and low-complexity regions using the NCBI standard and *Danio* species parameters, with output converted to GFF3 format using a custom script; 2. Known Repeats: Masking against known repeats from *Danio* species using the RepeatMasker database;3. De Novo Repeats: Masking using the custom repeat library generated by RepeatModeler. The results from all three masking runs were concatenated, and the final soft-masked genome FASTA file was generated using bedtools maskfasta with the combined GFF3 file. Gene annotation was performed by lifting over a robust annotation from the closely related reference genome. We used Liftoff (v1.6.3) to project the gene annotations from *Danio rerio* (GRCz11, NCBI GCF_000002035.6 GTF file) onto the newly assembled and scaffolded *D. albolineatus* genome. This approach leveraged the well-curated annotation of *D. rerio* to provide functional gene models for the novel assembly.

#### Bulk Segregant Analysis (BSA)

The genetic locus underlying the caudal fin color variation was mapped using a Bulk Segregant Analysis (BSA) approach comparing two distinct F2 pools with 23 specimens each derived from an original cross between the more Yellow-finned *D. albolineatus* and the more Red-finned *Danio roseus*. The two pools, designated “Yellow” and “Orange”, were sequenced to a mean coverage of 2x the number of specimens (see Appendix). Raw Illumina PE 150bp sequencing reads were quality-trimmed and filtered using fastp (v0.23.2). The cleaned paired-end reads were aligned to the final soft-masked *D. albolineatus* reference genome (RagTag-scaffolded with *D. rerio*) using Bowtie 2 (v2.4.5) with the --sensitive-local preset. The resulting alignments were filtered to remove unaligned reads and reads with low mapping quality (MAPQ < 20) using Samtools (v1.17). Duplicate reads were identified and removed using a series of steps involving samtools fixmate and samtools markdup to ensure unbiased variant calling. Single-nucleotide polymorphisms (SNPs) were called across all samples using BCFtools (v1.17) in a per-region, parallelized workflow. The mpileup and call commands were executed with the --multiallelic-caller option, annotating for allelic depth (AD) and total depth (DP). Variants were filtered based on quality (QUAL >= 20), mapping quality (MQ >= 20), and minimum per-sample read depth (MIN(FMT/DP) >= 5). Two distinct filtered VCF sets were generated: 1. Population Genetics Filter (PopGen): Used the base quality and depth filters defined above; 2. Association Filter: Used the PopGen filters and included an additional filter for minor allele frequency (MAF) to retain only common variants (MAF >= 0.05 and MAF <= 0.95). The association between genetic variants and the pooled phenotypes was assessed using two independent methods: 1. Allele Frequency Comparison: The PopGen VCF files were converted to Popoolation2 sync format for allele frequency comparison. Popoolation2’s (v1.201) fisher-test.pl script was employed to calculate Fisher’s Exact Test P-values for allele frequency differences between the Yellow and Orange pools at each SNP, implementing a dynamic coverage filter based on the interquartile range (IQR) of 10kbp window depth values (Q3 + 5 * IQR) to remove outliers; 2. Sliding Window Analysis: The Association VCF files were reformatted and analyzed using the QTLseqr R package (v2.1.0). This approach calculates the deltaSNP (difference in allele frequencies) and the G’ statistic (log-ratio of bulk allele frequencies, corrected for variation) in sliding windows of 1Mbp and 100kbp steps. Significant QTL regions were identified using a permutation-based null distribution (q-value <= 0.05). To identify the most robust candidate single-nucleotide polymorphisms (SNPs) underlying the caudal fin color, we implemented a three-tiered prioritization strategy: 1. Statistical Cross-Reference: Significant loci were defined by intersecting the outputs of both QTL mapping methods. Only SNPs falling within a genomic region identified by QTLseqr as significant (q-value <= 0.05) and having a Fisher’s Exact Test P-value greater than the 99.9th percentile threshold calculated by Popoolation2 were retained; 2. Parental Genotype Filtering: The resulting subset was further filtered using the fixed-genotype calls from the Grandparent samples. This stringent filter retained only SNPs that adhered to a model of Parental Fixed Difference (PfixedDiff), where the two grandparental lineages were fixed for opposing alleles (0/0 vs. 1/1). This step maximized the chance of identifying a true causal variant segregating in the F2 pools; 3. Functional Annotation: The final list of candidate SNPs was functionally annotated using SnpEff (v4.3t), employing the *D. rerio* lifted gene annotation to predict the potential molecular consequence (e.g., missense, nonsense, intron variant) of each variant. Nucleotide diversity (pi), Tajima’s D, and Weir and Cockerham’s Fst were calculated for the Yellow and Orange pools using grenedalf (v0.6.1) from the PopGen-filtered VCF set in non-overlapping 10kbp windows.

#### Behavioral assay

To test for a social aggregative behavioral preference for different phenotypes, wild-type *D. albolineatus*, red mutant *rgmb^vp67re1^* and yellow mutant *bdh1a^vp40ac1^* were used. All individuals were 1 year old and selected for comparable size and body shape. The behavioral arena consisted of a glass tank (51 × 28 × 25 cm) with bottom covered by small gravel to simulate a natural environment. Two identical glass chambers designated as male compartments (12 × 8 × 12 cm) were placed at opposite ends of the tank. These were connected by two transparent Plexiglas boards to create a female compartment (21 × 12 × 12 cm) that allowed the test female to swim freely between male chambers. The glass tank and all chambers were filled with system water to a depth of 12 cm. The arena was enclosed in a white-cloth-covered box (1 × 1 × 1 m) to diffuse light and minimize external visual disturbances. Full-spectrum LED lights (Draco Broadcast Dracast LED500, 5600 K) were positioned 1 m above the tank on both sides outside of the box to ensure uniform illumination. A front opening allowed the camera lens for video recording. All trials were recorded at 30 frames per second using a Panasonic HC-V770 digital camera. Two black dividers were placed on either side of the female compartment during setup. Two male fish were introduced into the male chambers and one female into the central compartment, each using separate fish nets. Male chambers were covered with transparent lids to prevent escape. Fish were acclimated for 10 min before testing. Testing began upon simultaneous removal of the dividers, and each assay lasted at least 5 min. After testing, fish were removed, and chambers were rinsed with RO water before the next trial. Each fish was used only once. To control for circadian effects on sensory behavior and motivation, all assays were performed between 08:30 and 11:30. The positions of the male genotypes were alternated between trials to avoid positional bias. Trials were excluded if the female showed no apparent interest in either male (i.e., swam continuously without spending more than 10 seconds near either male chamber). *N*=16 in wild-type vs. yellow trials (20 in total, 4 excluded); *N*=17 in red vs. yellow trials (20 in total, 3 excluded); *N*=14 in wild-type vs. red trials (17 in total, 3 excluded). The female’s horizontal distance from the chamber center was measured every 15 frames (0.5 s).

#### Parametric and non-parametric statistical analyses

Analyses of quantitative data were performed in JMP Pro 18 (SAS Institute, Cary NC), with transformations applied as noted in figure legends to correct for heterogenous variances in residuals as warranted.

## Notes

### Competing Interest Statement

The authors have declared no competing interest.

